# A basic framework governing splice-site choice in eukaryotes

**DOI:** 10.1101/2024.03.21.586179

**Authors:** Craig I Dent, Stefan Prodic, Aiswarya Balakrishnan, James Georges, Aaryan Chhabra, Sourav Mukherjee, Jordyn Coutts, Michael Gitonobel, Rucha D Sarwade, Joseph Rosenbluh, Mauro D’Amato, Partha P Das, Ya-Long Guo, Alexandre Fournier-Level, Richard Burke, Sridevi Sureshkumar, David Powell, Sureshkumar Balasubramanian

**Author notes:** These authors contributed equally – shared first authors (Craig I Dent, Stefan Prodic, Aiswarya Balakrishnan). These authors contributed equally – shared second authors (James Georges, Aaryan Chhabra). Max-Planck Institute for Plant Breeding and Genetics, Cologne, Germany (Craig I Dent). The Centre for Computational Biomedical Sciences, John Curtin School of Medical Research, Australian National University, Canberra, Australia (Stefan Prodic). University of Rochester Medical Center, University of Rochester, New York, USA. Author for correspondence Sureshkumar Balasubramanian School of Biological Sciences Monash University Clayton Campus VIC 3800 AUSTRALIA.

## Abstract

Changes in splicing are observed between cells, tissues, organs, individuals, and species. These changes can mediate phenotypic variation ranging from flowering time differences in plants to genetic diseases in humans. However, the genomic determinants of splicing variation are largely unknown. Here, we quantified the usage of individual splice-sites and uncover extensive variation between individuals (genotypes) in Arabidopsis, Drosophila and Humans. We used this robust quantitative measure as a phenotype and mapped variation in splice-site usage using Genome-Wide Association Studies (GWAS). By carrying out more than 130,000 GWAS with splice-site usage phenotypes, we reveal genetic variants associated with differential usage of specific splice-sites. Our analysis conclusively shows that most of the common, genetically controlled variation in splicing is *cis* and there are no major *trans* hotspots in any of the three analyzed species. High-resolution mapping allowed us to determine genome-wide patterns that govern splice-site choice. We reveal that the variability in the intronic hexamer sequence (GT[N]_4_ or [N]_4_AG) differentiates intrinsic splice-site strength and is among the primary determinants of splice-site choice. Experimental analysis validates the primary role for intronic hexamer sequences in conferring splice-site decisions. Transcriptome analyses in diverse species across the tree of life reveals that hexamer rankings explains splice-site choices from yeast to plants to humans, forming the basic framework of the splicing code in eukaryotes.

## Introduction

RNA splicing is a critical gene regulatory process in which specific regions of the pre-mRNA (introns) are removed with the joining of the adjacent regions (exons) to produce the mature mRNA that encodes the protein. RNA splicing affects growth, development, and response to external stimuli (Marasco and Kornblihtt, 2023; Rogalska et al., 2023). Changes in splicing can either alter the protein produced or can trigger nonsense-mediated mRNA decay (NMD), regulating the amount of functional protein (Nasif et al., 2018). RNA splicing can differ between cells, tissues, organs, genotypes, and species (Marasco and Kornblihtt, 2023) and in response to developmental and environmental cues, shaping phenotypes with possible evolutionary consequences (Ule and Blencowe, 2019). Changes in RNA splicing are also observed in various diseases, including cancer (Bradley and Anczukow, 2023; Rahman et al., 2020) making RNA splicing a potential avenue for therapeutics. Thus, there is an interest in understanding the mechanisms through which genetic variation can affect RNA splicing and disease (Rogalska et al., 2023).

A vast majority of introns contain a consensus GU at their 5’ end (donor) and an AG at their 3’ end (acceptor), with a small proportion of introns harbouring alternative motifs (5’ GC or AT and 3’ AC) (Lerner et al., 1980; Parker et al., 2022; Rogers and Wall, 1980; Wong et al., 2018). In addition, a branch point “adenosine (A)” and a polypyrimidine stretch towards the 3’ end of introns are known sequence features required for splicing (Aebi et al., 1986; Petrillo, 2023; Rogalska et al., 2023; Smith et al., 1993; Wahl et al., 2009; Zarnack et al., 2020). However, in an RNA molecule, there are multiple GU/AG sequences that could be used in a condition- or genotype-dependent manner. The rules that govern which GU/AG become splice-sites is still unclear. Massively parallel splicing assays have been performed to assess the impacts of sequence variation in consensus sequences (Roca et al., 2012; Rosenberg et al., 2015; Wong et al., 2018). While consensus sequences are known (Lee and Rio, 2015; Roca et al., 2012; Rogalska et al., 2023; Wong et al., 2018), how specific sequence combinations within the consensus affect splice-site choice is still unclear.

Since sequence variation can affect splicing, there is considerable interest in predicting the impact of genetic variation on splicing. Programs such as SpliceAI, Pangolin, SpliceVault, SPLAM and other such approaches use various artificial intelligence (AI) methods to accurately predict the impact of genetic variation on splicing (Chao et al., 2023; Dawes et al., 2023; Jaganathan et al., 2019; Zeng and Li, 2022; Zhang et al., 2023; Zhang et al., 2022). However, it is often difficult to comprehend the biological parameters through which AI-based approaches achieve that high predictive power. There are efforts to understand the logic through which these approaches succeed (Liao et al., 2023). Further, the performance of models trained on one species is low when applied to other species (Chao et al., 2023), which suggests that the genetic differences in splice-site choice are not well understood.

Accurate measurement of phenotypes is critical for successful and trustworthy genetic mapping. However, efforts to map splicing variation suffers from the accuracy of the splicing phenotypes used in the analysis. For example, splicing QTL (sQTL) analysis has been conducted on diverse datasets (Atla et al., 2022; Consortium, 2020; Garrido-Martin et al., 2021; Jin et al., 2023; Khokhar et al., 2019; Li et al., 2018; Tian et al., 2022; Walker et al., 2019), but the quantification of splicing often suffers from a lack of specific focus on individual splice-sites, which prevents deciphering a direct association between the genetic variant and a specific splice-site (Consortium, 2020; Dent et al., 2021; Khokhar et al., 2019). In addition, the sQTL approach often tests only SNPs within a region around the genes as opposed to genome-wide association, typically increasing the risk of spurious associations (Consortium, 2020; Khokhar et al., 2019; Tian et al., 2022). Therefore, an accurate quantification of individual splice-site usage coupled with GWAS can help map associated genetic variants, which will be highly valuable in deciphering genomic features of the splicing code.

We have previously defined a measure of splicing that focuses on the usage of individual splice-sites (Dent et al., 2021) as opposed to more general features such as splicing events, isoforms, exon inclusions, intron clusters, or localized splice graphs (Dent et al., 2021; Li et al., 2018; Schafer et al., 2015; Shen et al., 2014; Vaquero-Garcia et al., 2016; Wang et al., 2008). This measure of splice-site strength/usage was defined by Reed and Maniatis as the ability of the splice-site to participate in a splicing reaction (Reed and Maniatis, 1986). It can be accurately measured with SpliSER (Splice-site Strength Estimate from RNA-seq) and this quantitative measurement can be used as a phenotype in GWAS to detect robust associations (Dent et al., 2021).

Here, we quantified individual splice-site usage leveraging RNA seq data from: i) the 1001 Genomes Project from *Arabidopsis thaliana* (Consortium., 2016; Kawakatsu et al., 2016), ii) male and female *Drosophila melanogaster* lines from the DGRP (Huang et al., 2015; Huang et al., 2014), and iii) human heart tissue from the GTEx project (Consortium, 2013; Siminoff et al., 2018). We show that there is extensive genetically controlled variation in individual splice-site strengths. By carrying out more than 130,000 GWAS using these data, we demonstrate that most of the variation in splice-site strength/usage is driven by *“cis”* regulatory variation. With high-resolution GWAS that map around splice-sites, we infer specific nucleotides for each position that are best suited to promote splicing and identify features that govern splice-site choice. We infer and show that intronic hexamers encompassing the splice donor and acceptor sites are the primary determinants of splice-site choice. Using average splice-site strengths, we develop a rank order of hexamers and show that most splice-site choices in the transcriptomes of diverse species can be explained by hexamer rankings. Our findings suggest that hexamer ranking forms the basic logical framework that explains splice-site choice across eukaryotic organisms.

## Results

### Extensive variation in splice-site usage in Arabidopsis, Drosophila, and humans

To assess splicing variation, we first quantified the usage of all splice-sites from Arabidopsis (Kawakatsu et al., 2016), Drosophila (Mackay et al., 2012) and Humans (Consortium, 2013) with SpliSER (Dent et al., 2021) and obtained splice-site usage data for a total of 767,199 splice-sites representing 39,734 genes from three different species (Table 1). Next, we computed: a) the range of splice-site usage (difference between the highest and lowest SSE) in individuals, and b) the variance in the usage of every splice-site. We observed extensive variation in the usage of individual splice-sites. Almost three-quarters of the human sites displayed more than 20% difference in their usage between individuals, while a half (50%) and a quarter (25%) of the sites showed such differences in Drosophila and Arabidopsis, respectively (Fig. 1A & D, Supplementary Figure S1A, Supplementary Table S1). In summary, we observed 430,345 splice-sites (56.1%) out of a total of 767,199 sites that displayed more than 20% difference in their usage between extreme individuals (Supplementary Table S1). The transcriptomes from Arabidopsis (Kawakatsu et al., 2016) and Drosophila (Huang et al., 2014) had replicates, which we exploited to calculate broad-sense heritability of splice-site usage and found that a substantial number of sites (>25%) displayed high heritability (>0.3 in Arabidopsis and even higher in Drosophila) in their strength. This suggested that there is a substantial genetic contribution to splicing variation that could be mapped. We selected all splice-sites for which there are at least 100 observations and that are in the upper quartiles of both variance and heritability, which resulted in a total of 17,140 sites in Arabidopsis, 7,711 sites in the Drosophila female and 8,003 in Drosophila male datasets. Human transcriptome data lacked replicates and therefore, we took all 97,796 sites that had data for at least 100 individuals and fell in the upper quartile of variance. In total, we took splice-site strength data from 130,650 sites for subsequent GWAS (SpliSER-GWAS).

**Table 1.** A summary of the sites analyzed in Arabidopsis, Drosophila, and humans. Assessable sites refer to the number of sites for which SSE values could be calculated from at least 100 individuals.

| Species | Total splice-sites | Corresponding number of genes | Assessable splice-sites | Number genes |
| --- | --- | --- | --- | --- |
| *Arabidopsis thaliana* | 219,336 | 15,099 | 112,729 | 9,804 |
| *Drosophila melanogaster* (Males) | 92,119 | 9,902 | 73,130 | 8,114 |
| *Drosophila melanogaster* (Females) | 76,626 | 8,197 | 65,551 | 7,215 |
| Human heart atrial tissue | 455,744 | 14,733 | 391,650 | 12,658 |

**Figure 1.**
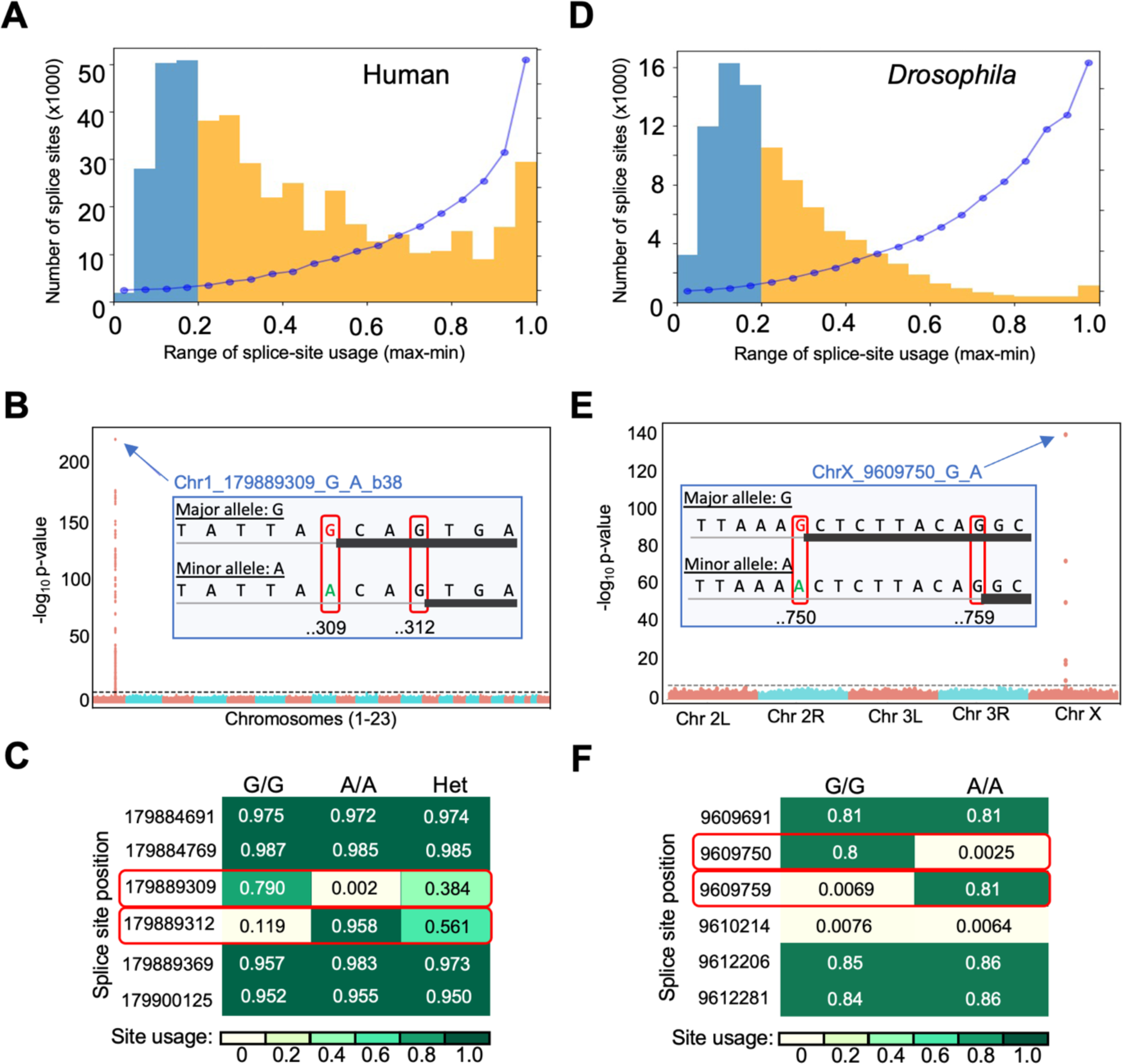
Splice-site usage varies extensively between individuals and can be mapped accurately with SpliSER-GWAS. A) Distribution of splice-sites that show differences in their usage between individuals in humans. The sites are grouped based on the maximum difference in their usage between individuals. Yellow region highlights the sites that display more than 20% difference (0.2 difference in) between extreme samples. Blue dots represent the average variance in splice-site usage between all individuals for each of the bins. B) Manhattan plot of the splice-site mutation at chr1:179,889,309. SpliSER-GWAS analysis identifies causal SNP for variation in the usage of splice-sites at the *TOR1AIP1* locus in humans. A schematic of the sequences surrounding two competing splice-sites in *TOR1AIP1* is shown. The splice-site mutation at chr1:179,889,309 (309) allows the usage of chr1:179,889,312 (312) as a splice-site and variation in the usage of both sites map to the 309 polymorphism. C) Average splice-site usage of the sites 309, 312 and the neighboring splice-sites. The effect of the mutation at 309 is specific to 309 and 312 and does not affect neighboring sites. D) Distribution of splice-sites that show differences in their usage between individuals in Drosophila. E) Manhattan plot of the for the splice-site mutation at in the Drosophila *HECW* gene. SpliSER-GWAS analysis identifies causal SNP for variation in the usage of splice-sites at the *HECW* locus in Drosophila. A schematic of the sequences surrounding two completing splice-sites at the *HECW* gene. A mutation in the *HECW* gene (Fbgn0261931-CG42797) of Drosophila at position chrX:9,609,750 (750) abolishes a splice acceptor site that leads to an increased usage of a site at chrX:9,609,759 (759). F) Average splice-site usage of the sites 750 and 759 and the neighboring splice-sites. The effect of mutation at 750 is specific to 750 and 759 and does not affect neighboring sites.

### SpliSER-GWAS is specific and captures causal variants for differential splice-site usage

SpliSER quantifications are accurate and specific (Dent et al., 2021) and can be validated experimentally (Supplementary Figure S2). To assess the sensitivity of our approach to identify causal variants, we took advantage of mutations at splice-sites, which will have a causal impact on splice-site usage and must be reflected in GWAS. Such mutations often lead to usage of an alternative site. We reasoned that we should identify the splice-site mutation as the highest associated SNP for both the mutated site and the alternative site. For example, at the *TOR1AIP1* locus in humans, there are individuals with a G to A mutation at the splice acceptor site at chr1:179,889,309. In individuals with this mutation, another acceptor site at chr1:179,889,312 is used (Fig 1B). We identified the G to A mutation at chr1:179,889,309 as the highest associated SNP for variation in the usage of both sites (chr1:179,889,309 & chr1:179,889,312) (Fig 1B), while the other sites on the same gene were unaffected (Fig 1C). We observed similar examples in Drosophila (Fig 1E, F) and Arabidopsis (Supplementary Figure S1B-D), which confirmed that SpliSER-GWAS is specific, accurate and in principle can identify causal SNPs for differential splice-site choice.

To assess the extent to which we can capture such variation, we analyzed all splice-site mutations that occur in at least in 5% of the individuals (minor allele frequency ≥0.05) by GWAS. This resulted in a total of 58 sites in Arabidopsis, 63 and 34 sites in Drosophila males and females, respectively, and 48 sites in the Humans (Supplementary Table S2). 152 of these 203 sites gave clean GWAS peaks (∼75%; 43/58 in Arabidopsis, 53/63 in Drosophila males, 27/34 in Drosophila females and 29/48 in Humans). More importantly, we identified the splice-site mutation as the highest and/or closest associated SNP in 34/58 sites (59%) in Arabidopsis and 29/48 sites (60%) in human heart tissue, 36/63 (57%) in Drosophila males and 16/34 (47%) in Drosophila females. These findings suggest that SpliSER-GWAS can capture up to ∼60% of the mappable causal variants for large-effect.

### Most of the genetically associated splicing variation is *“cis”*

From 130,650 splice-site usage phenotypes, we detected associations for a total of 21,988 sites (4,951 in Arabidopsis, 7328 in Drosophila and 9709 in Human; Supplementary Tables S3-S5). To assess the genomic architecture of splicing variation, we plotted the positions of splice-sites against their highest associated variants in all three species (Fig 2, Supplementary Figure S3, S4). We observed that most of the determinants of splicing variation mapped within 1Mb from the splice-site in all three tested species (Supplementary Tables S3-S5). We called these as ‘*cis’* and the rest as ‘*trans’*. *Cis-*associations represented 78 to 93% of all associations (Table 2), confirming that most genotype-dependent splicing variation is mediated via *cis-* rather than *trans-*genetic variation in Arabidopsis, Drosophila, and humans. While we do map verifiable *“trans”* associations (Supplementary Figure S2B), we did not observe any major *“trans”* hot spots (Fig 2, Supplementary Figures S3-S4).

**Figure 2.**
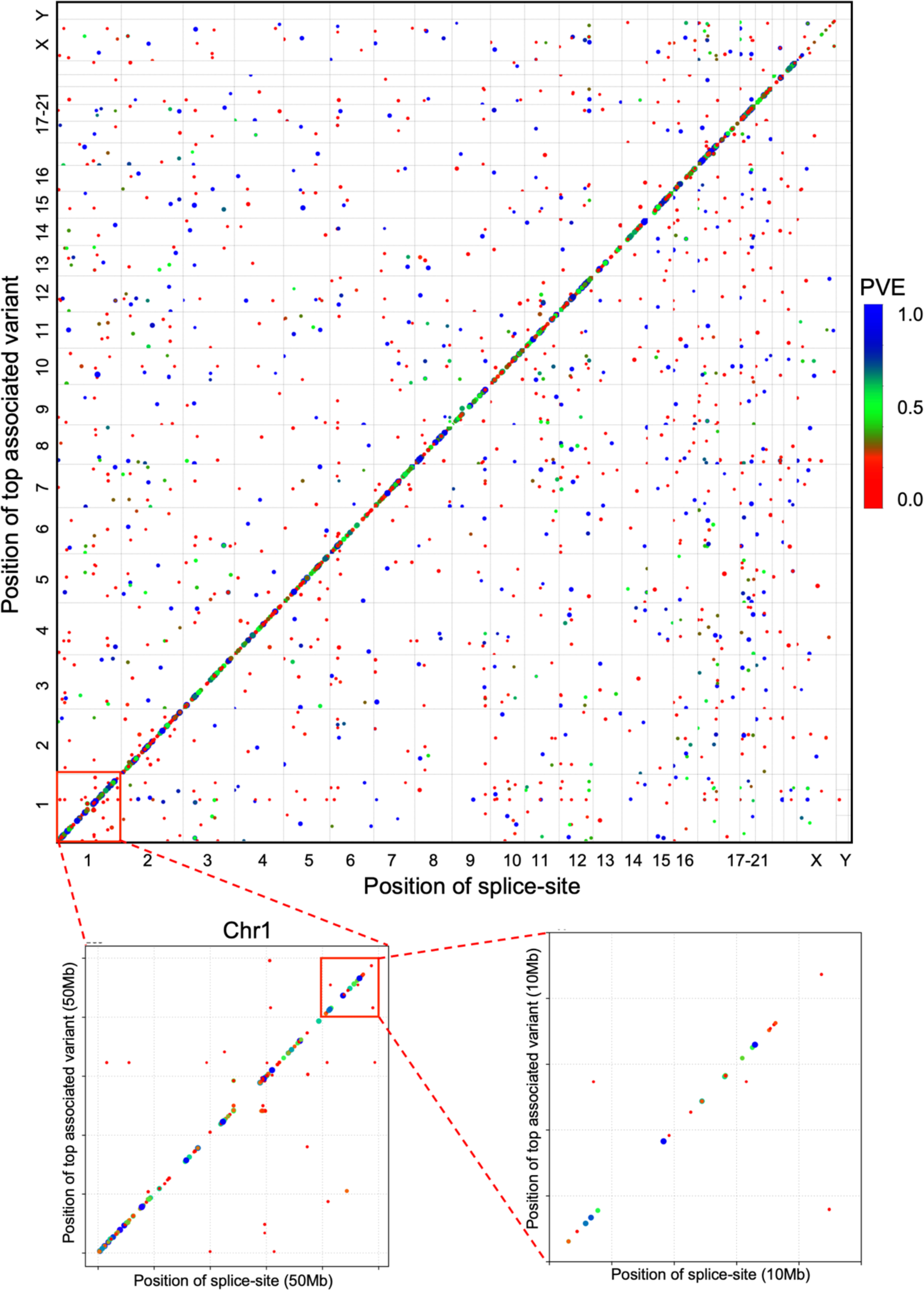
Splicing variability is mostly “*cis”* regulated. A-C) Scatter plot of splice-site positions and their highest associated SNPs in the human heart across the human genome. Chromosome 1 is zoomed in at two different levels to show the *cis* nature of associations. Colour scale represent the percentage of variance (PVE) explained by the associated top SNP. The sizes of the dots are also correlated with PVE.

**Table 2.** A summary of associations detected in different RNA-seq datasets.

| Species/<br>Phenotype | Total<br>associations | Number of “*cis*”<br>associations | Number of “*trans*”<br>associations | Percentage of “*cis*”<br>associations |
| --- | --- | --- | --- | --- |
| Arabidopsis | 5277 | 4361 | 916 | 82.6% |
| Drosophila–a -<br>males | 3897 | 3559 | 338 | 91.3% |
| Drosophila–a -<br>females | 3453 | 3143 | 310 | 91% |
| Human heart<br>atrial tissue | 9872 | 7711 | 2161 | 78.1% |

### Intronic regions drive most of the variation in splice-site usage

To assess the features of splicing variation, we plotted the distribution of distances of the highest and closest-associated SNPs to the splice-site for all sites in all three species, which revealed that most of the associations map in and around the splice-site (Fig 3A, B, Supplementary Figure S5A-B). In all our position and distance calculations, the splice-site G (G of GT as well as G of AG) is given the position 0 and the downstream and upstream positions were given positive and negative coordinates, respectively. The same notion is followed throughout this manuscript in all analysis. In fact, we observed 32% (1410/4361, 611 donors and 799 acceptors) associations in Arabidopsis where the highest-associated SNP fell within 100bp of the splice-site; we saw 24% in Drosophila (1771/7350, 772 donors and 999 acceptors) and 9% (651/7711, 281 donors and 370 acceptors) in humans. These numbers almost doubled if we considered the SNPs within the peak closest to the site (60%, 52% and 23% in Arabidopsis, Drosophila, and humans, respectively). To assess whether intronic or exonic nucleotides make the biggest difference to splice-site strength, we asked how much of the variation in splice-site strength/usage across the genome can be explained by individual nucleotides surrounding the splice-site by taking all splice-sites into account. This analysis suggests that splice-site strength is mostly affected by intronic rather than exonic sequences (Supplementary Figure S6).

**Figure 3.**
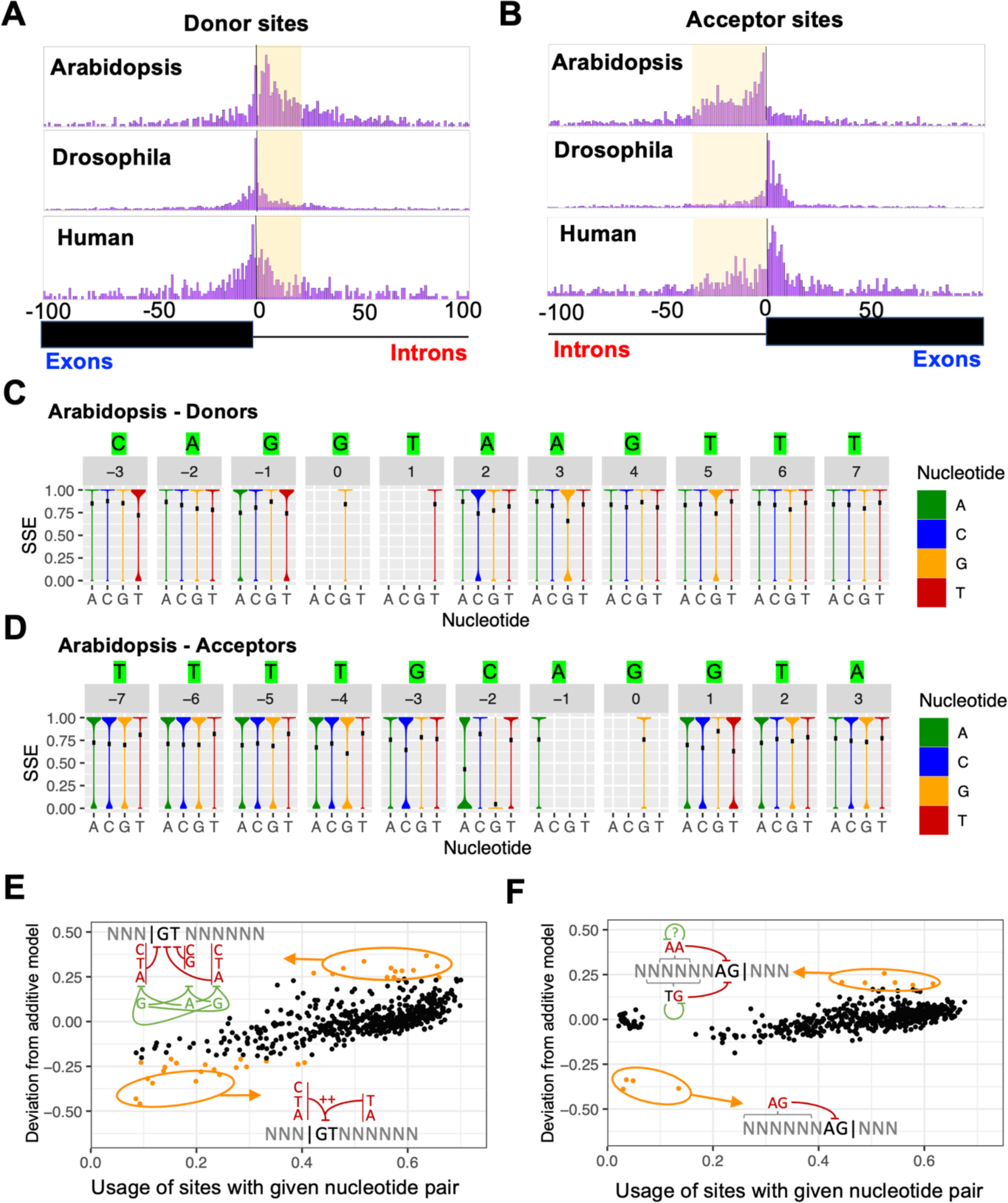
Genetic variation affecting splice-site choice is often near the splice-site and high resolution SpliSER-GWAS allows inferring best nucleotides that promote splicing. A-B) Distribution of the distances of the highest associated SNPs detected in SpliSER-GWAS for acceptors (A) and donors (B) in Arabidopsis, Drosophila and Humans. The “G” of “GT” or “AG” is plotted as position 0. Intronic regions are shaded for clarity. C-D) Violin plots depicting the mean splice-site strength estimate (SSE) for all splice-sites harbouring each of the four possible nucleotides around the splice-sites. For acceptors (C), positions -7 to +3 is and for donors (D), positions -3 to +7 is shown. The nucleotide with the highest mean splice-site strength at each position is highlighted on the top. E-F) Pairwise combinations of nucleotides around the splice site whose Splice-site Strength Estimate (SSE) deviate from a naïve additive model. (E) – Donor sites (GT only), (F) - Acceptor sites (AG only). Each point represents a single pairwise combination of nucleotides around the splice site, -3 to +7 in donors, and -7 to +3 in acceptors. Nucleotide pairs are plotted with the SSE (x-axis) and deveiation from the additive model (y-axis). The pairs which deviate from the additive model are marked in orange.).

### Specific nucleotides in and around splice-sites differentiate splice-site strengths

To assess genetic variation that drives splice-site strength, we looked at the distributions of the distances of the highest-associated SNPs from their respective splice-sites (Fig 3A, B). A substantial fraction of the variants mapped near splice-sites. We observed an intronic bias in Arabidopsis, but the pattern was more diffused in Drosophila and Humans, which suggested that the genetic architecture of splicing variation differs slightly between species (Fig 3A, B). In each of the associations from GWAS, there are SNPs (alleles) associated with increased (splice-promoting) or decreased (splice-reducing) usage of the splice-site. We inferred the most frequent bases among the splice-promoting (best) and splice-reducing (worst) nucleotides for each position (Fig 3C, D, Supplementary Figure S7A-C, Supplementary Table S6). These sequences differed in general indicating that these patterns are not driven purely by background nucleotide frequencies.

To assess the functional impact, we designed best and worst synthetic introns containing the splice-promoting (best intron) and splice-reducing (worst intron) nucleotides at each position (Supplementary Table S7). We introduced these synthetic introns into the coding region of mCHERRY between AG and GT (AG]-[GTintronAG]-[GT). Since the inferred “worst” sequences also lacked the consensus GT donor and AG acceptor sites, we modified the intron to include them. We noticed that there were adenosines at positions -19, -25 and -31 that could be potential branch points. We transfected these constructs into HEK293T cells. Consistent with our expectations, we observed strong fluorescence in cells that were transformed with the construct having the best intron (Supplementary Figure S8A, S8B). Cells that were transformed with the worst intron (despite harboring GT-AG & branch point) displayed no fluorescence. RT-PCR analysis confirmed the appropriate splicing of the best intron and no splicing of the worst intron, consistent with our prediction (Supplementary Figure S8A, S8B). We conclude that allelic variations beyond the GT/AG and the branchpoint adenosine have a significant impact on splicing and contribute to splicing variation.

We also calculated the average strength of all splice-sites across the genome for every nucleotide at every position in and around the splice-sites, which also provided a measure of the best base for each position surrounding the splice-site (Fig 3C, D, Supplementary Figure S9). This analysis revealed nucleotides that significantly affect splice-site strength. In splice donor sites, in Humans and *Drosophila* the most deleterious effects were seen with the inclusion pyrimidines 2-4 bp downstream of the splice-site (Supplementary Figure S9, Supplementary Table S8). Further, in *Drosophila*, the absence of a +4 Guanine accounted for a 29% change in splice site usage, based on its presence or absence (Supplementary Figure S9, Supplementary Table S8). In *Arabidopsis,* the pattern was different; the largest difference was due to the absence of a Guanine at the -1 position (12% difference; Fig 3C, Supplementary Figure S9, Supplementary Table S8), where the G at -1 position is almost invariant. At splice acceptor sites, we saw the largest deleterious effect coming from purines upstream of the splice-site. This impact was less pronounced in Arabidopsis, where sensitivity to purines at the -2 position could be observed. In all species, the largest single-nucleotide impact on splicing was the negative impact of Guanine at position -2 (Supplementary Figure S9, Supplementary Table S9).

### Combinatorial effects of nucleotides in and around splice-sites

Pairwise interactions between nucleotides have been previously analyzed through massively parallel splicing assays in the context of three genes (Wong et al., 2018). Using genome-wide splice-site strength data, we looked for non-additive pairwise interactions between nucleotides at different positions around the splice site. In acceptor sites, we consistently saw that an AG dinucleotide upstream of the measured site had a larger decrease in splice site usage than would be expected by an additive interaction of either an A or G from those positions (Fig 3E-F). This was most prominent at positions -7 through -4, with splice-site usage being reduced by 25-60% more than expected (Fig 3E-F, Supplementary Figure S10, Supplementary Table S10-S11). We also observed this pattern downstream of the splice site, but to a lesser degree (Supplementary Figure S10, Supplementary Table S10-S11). We also observed several other pairwise interactions that suggested that specific combinations of sequences have significant impacts on splice-site strength (Fig 3E-F, Supplementary Figure S10-S13, Supplementary Table S10-S11). The multi-layered interactions of nucleotides around the splice site motivated us to ask which regions around the splice site could best capture the variation in splice site usage.

### GT[N]_4_ and [N]_4_ AG hexamer sequences are the primary determinants of splice-site choice

Given that intronic region primarily accounts for variation in splice-site usage, we considered which regions of the intron could account for variation in the intrinsic strength of splice-sites and potentially comparable across species. We considered three aspects. First, we checked which length *k*-mers are present in most possible sequence combinations at splice-sites. For example, a tetramer (e.g., GTNN) has 16 possible sequence combinations and all 16 could be found at the splice-sites. However, an octamer (GT [N]_6_) has 4096 sequence combinations, but only some of these could be found at splice-sites. Therefore, a substantial number of octamers are not comparable across species. Second, we assessed the ability of *k*-mers to differentiate GT/AGs around splice-sites. The longer the *k-*mer, the more likely it can uniquely differentiate a splice-site among competing sites. Third, we grouped all possible splice-sites into distinct *k*-mer groups, used their splice-site strengths to calculate the average strength of that *k*-mer and generated *k-* mer rankings. We systematically scanned a 100bp window (50bp upstream and downstream) surrounding every individual splice-site for all the potential competing GT or AG dinucleotides, extracted corresponding *k*-mers (GT, GTN, GTNN etc.) and queried in what proportion of splice-sites *k*-mer ranking can explain splice-site choice. We determined the percentage of splice-site choices explained by different *k*-mer lengths (i.e., the percentage of sites in which the highest-ranked *k*-mer being utilized as the splice site) and found that GT[N]_4_ hexamers explain most of the donor-site choices (Table 3). For acceptor sites, we found both hexamers [N]_4_AG as well as [N]_5_AG had similar scores, through the hexamers on an average slightly outperformed the heptamers across species (Table 3). All three species gave similar patterns (Supplementary Table S12). We conclude that hexamers present an optimal *k*-mer for cross-species comparisons to explain splice-site choice. In addition, we observed significant correlations of the frequency of hexamers with splice-site strength but not with their frequency in gene bodies (or genome, Supplementary Figure S14), which indicates that there is natural selection favoring different hexamer sequences at the splice-site, potentially due to their functional impact. In summary, hexamer ranking explained most splice-site choices across the transcriptomes of eukaryotes from plants to humans suggestive of a fundamental basic framework.

**Table 3.** Percentage of splice-site choices explained in three different species with different length intronic *k*-mers.

| Sequence | *Arabidopsis* | *Drosophila* | Humans | Average |
| --- | --- | --- | --- | --- |
| **Donors** |  |  |  |  |
| GT | 0.66 | 0.76 | 0.87 | 0.76 |
| GTN | 28.56 | 15.02 | 26.73 | 23.43 |
| GTNN | 38.45 | 48.73 | 52.42 | 46.53 |
| GTNNN | 56.06 | 77.47 | 69.87 | 67.80 |
| **GTNNNN** | **58.26** | **81.41** | **72.86** | **70.84** |
| GTNNNNN | 48.95 | 65.84 | 69.05 | 61.28 |
| GTNNNNNN | 24.60 | 40.90 | 53.24 | 39.58 |
| **Acceptors** |  |  |  |  |
| AG | 0 | 1.93 | 0 | 0.64 |
| NAG | 30.24 | 28.46 | 19.49 | 65.19 |
| NNAG | 48.95 | 46.04 | 32.14 | 42.37 |
| NNNAG | 60.98 | 64.05 | 46.46 | 57.16 |
| **NNNNAG** | **65.73** | **71.02** | **55.25** | **64.00** |
| NNNNNAG | 65.71 | 65.03 | **57.72** | 62.00 |
| NNNNNNAG | 53.92 | 43.29 | 51.10 | 49.43 |

### Hexamer ranking explains mutational impacts on splice-site choice in diverse species

Since intronic hexamers explained splice-site choices, we tested whether they could explain the effects of previously described mutations. Two different transcripts are produced from the *FLOWERING LOCUS M (FLM)* gene of Arabidopsis based on the splicing of the donor GT of the first intron with AG1 or AG2 acceptor sites. Generally, AG1 is preferred over AG2 (Capovilla et al., 2017; Lutz et al., 2015; Sureshkumar et al., 2016). Hexamer analysis revealed that AG1 harbors a stronger hexamer compared with AG2, which can explain its preference (Fig 4A). In addition, a mutation (GG to AG at position chr1:28,958,437) in the second intron of *FLM*, abolishes of the use of either of these partner sites (Hanemian et al., 2020). It turns out that the mutation creates a new AG acceptor site with the strongest hexamer of the three, which can explain the abolition of the use of AG1/AG2 (Fig 4A).

**Figure 4.**
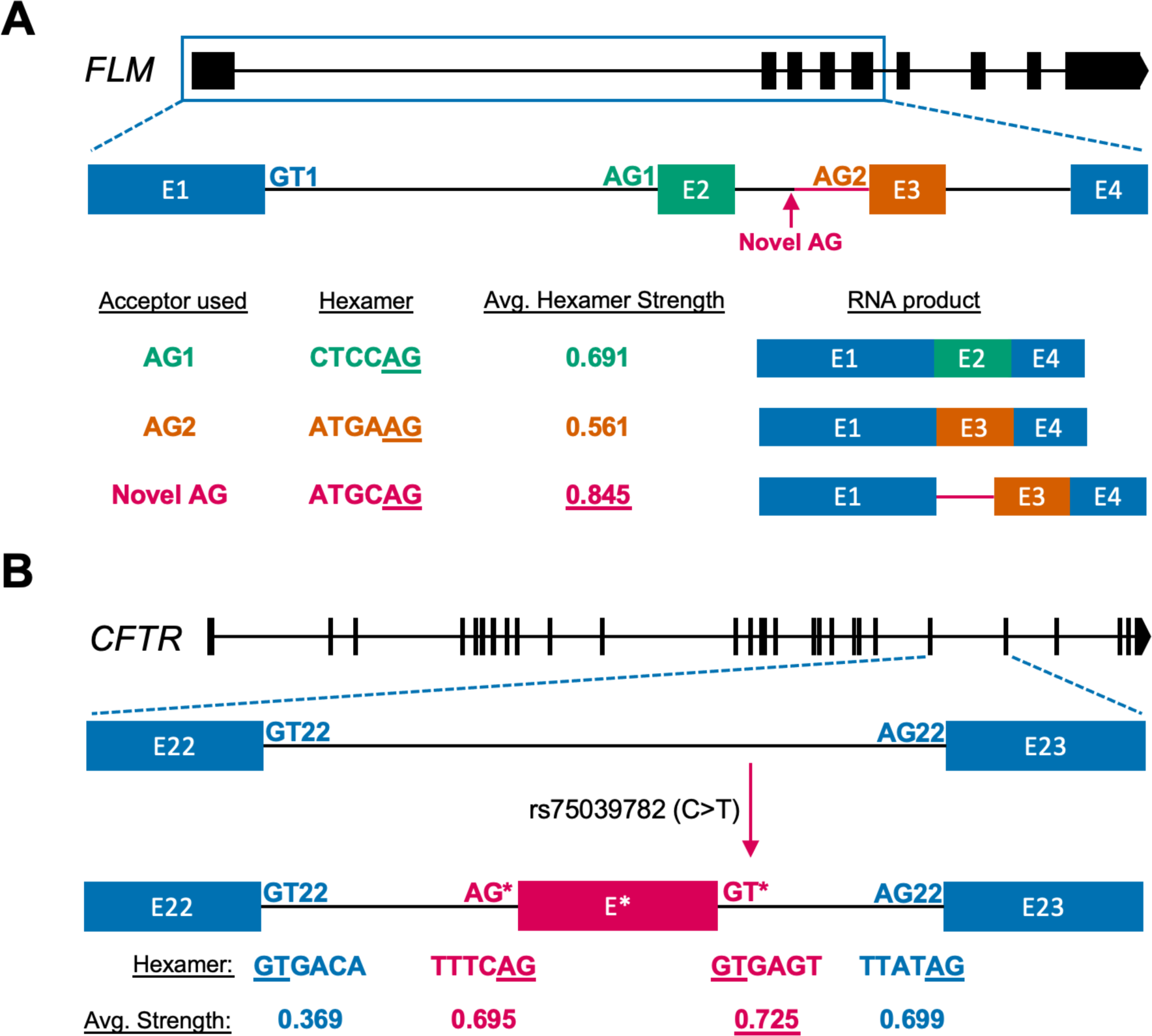
Hexamer ranking explains mutational impacts on splice-site choice. A) A schematic representing the natural mutation that creates a competing AG at the *FLM* locus. RNA products arising out of the three competing AGs are shown along with their average splice-site strength. Hexamer ranking explains why AG1 is preferred over AG2 and why the new AG outcompetes both. B) Hexamer ranking explains why rs75039752 SNP at the CFTR gene leads to inclusion of a pseudo exon and leads to cystic fibrosis. rs75039752 introduces a new splice-site with the hexamer of higher rank that outcompetes the normal donor, which hijacks the natural partner. This allows another acceptor, which is not normally used to be used now to include a pseudo exon, resulting in a frameshift and an impaired CFTR protein.

Similarly, in humans, several mutations in the *CFTR* gene are associated with splicing differences (Deletang and Taulan-Cadars, 2022). One of the common mutations is *rs75039782* (C to T) in the 22^nd^ intron of the *CFTR* gene, which generates a new donor GT site (Augarten et al., 1993). This mutation leads to a pseudo-exon containing a premature stop codon, resulting in CFTR deficiency and cystic fibrosis. Hexamer analysis explains this mutational impact: The *rs75039782* mutation generates a donor site with a stronger hexamer than the canonical donor site, resulting in an otherwise unused upstream acceptor site in the intron now being used in partnership with the canonical donor site that leads to the inclusion of the pseudo-exon (Fig 4B).

### Hexamer rank order-driven splice-site choice is conserved from yeast to plants to humans

To assess whether the hexamer rank order can explain splice-site choices across eukaryotes, we analyzed publicly available RNA-seq data to compute hexamer ranks based on their strength and frequency across 25 eukaryotic species and asked what proportion of the splice-site choice could be accounted for by hexamer rankings. Indeed, hexamer ranking based on splice-site strength explained most splice-site choices (∼60-85%) in diverse species (Supplementary Table S12-S14). Next, we computed pairwise rank-correlations of hexamer rankings based on either strength or frequency and constructed dendrograms based on their R^2^ values. R^2^ values in all species comparisons were positive, and reflected phylogenies (Supplementary Table S15, Supplementary Figure S15A).

Such a congruence between of the dendrograms and the phylogeny suggested a potential basic feature governing the biology of splice-site choices. One obvious candidate for donor sites is the U1snRNA that base pairing with the 5’ splice-sites. The most frequent and the strongest hexamer GTAAGT (Supplementary Table S13) is complementary to the most common U1 snRNA base pairing site of CAUUCA. Therefore, we grouped the hexamers using sequence distances from the U1 snRNA binding site (GTAAGT) and asked whether the average strengths of these hexamer groups reflect sequence distance from the U1 snRNA binding site. Sequence distance matrices do not provide the same level of resolution as the average strengths. Nevertheless, we observed a near perfect correlation between the sequence distances and the hexamer rankings in Arabidopsis, Drosophila and Humans (Supplementary Figure S15B). This data suggests that, at least for the donor sites, hexamer-based differences in splice-site choices is primarily driven by U1 snRNA base-pairing properties. For the 3’ acceptor sites, we computed the distances from the hexamer with the highest strength (TTGCAG) and found a very similar correlation between sequence distances and hexamer rankings (Supplementary Figure S15C). Taken together, these results indicate that intronic hexamer sequences form a basic feature of the splicing code in eukaryotes.

### Testing of hexamer sequences as the determinants of splice-site choice

We took two different approaches to verify whether hexamer rankings are the primary determinants of splice-site choice. First, we leveraged data published by Rosenberg et al and carried out a meta-analysis with a focus on hexamer ranking (Rosenberg et al., 2015). Rosenberg et al transfected human cells with a multitude of mini-gene constructs with embedded random sequences that can form novel splice-sites and analyzed resulting constructs (DNA) and transcripts (RNA) by sequencing. This dataset allowed the direct testing of thousands of competing hexamers in cell culture experiments. We ran SpliSER on this RNA-seq data and quantified the splice-site usage of all splice-sites. A total of 466,000 competing donors allowed us to evaluate ∼4,000 unique pairs of competing hexamers. We queried how many of the winning splice-sites could be explained by the hexamer rankings and found hexamer ranking could explain 66% of splice-site choices (Fig 5A). This percentage increased to 91%, when observing constructs where the differences in hexamer strengths were more than 25%, providing further evidence that hexamers are among the primary determinants of splice-site choice (Fig 5A).

**Figure 5.**
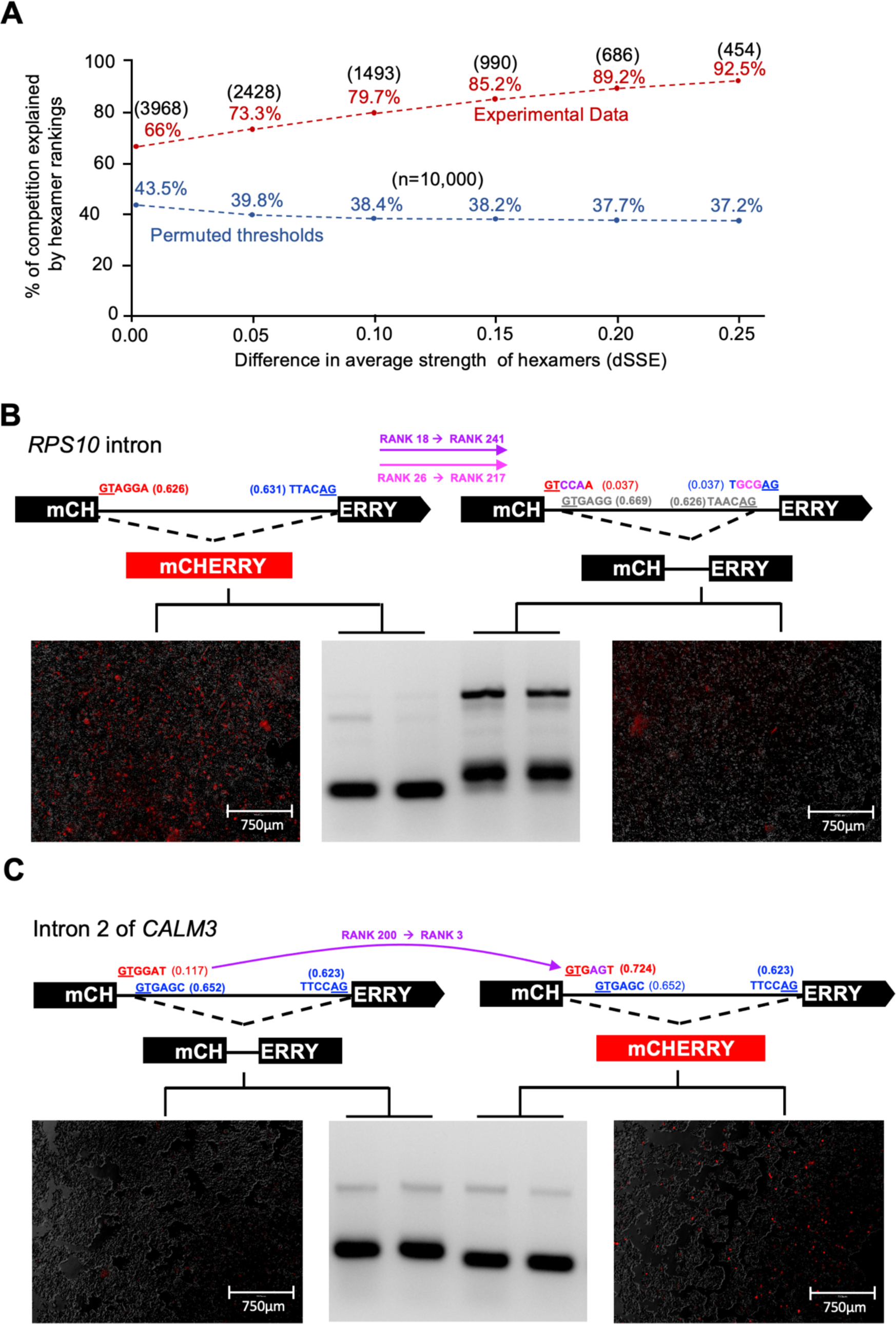
Hexamer rankings determine splice-site choice in experiments. A) Direct comparison of competing hexamers in minigene constructs of Rosenberg et al. Blue line represents permuted thresholds calculated from 10,000 permutations and the actual experimental data (red) with varying combinations of hexamers. The number of unique competing pairs tested is shown above the percentages. B, C) Conversion of a good intron into a bad intron (B) or bad intron into a good intron (C) through change of hexamers. mCHERRY fluorescence is shown along with the RT-PCR amplification.

To further assess this idea, we designed three types of mini-gene constructs with the mCHERRY reporter gene. First, we scanned for human introns that are removed with more than 95% efficiency (i.e., donor and acceptor SSE ≥0.95, good intron) or less than 10% efficiency (i.e., donor, and acceptor SSE ≤0.1, bad intron) across all individuals in the GTEx dataset. We reasoned that by changing the hexamers, we can convert bad introns into good ones and *vice versa.* An intron in the *RPS10* gene with splice donor site at chr6:34,425,071 and acceptor site at chr6:34,424,840 is removed with almost 100% efficiency (>99% across all GTEx individuals). We generated constructs where the hexamer sequences at the donor and acceptor sites (GTAGGA [Rank 18, SSE-0.626] / TTACAG [Rank 26, SSE-0.631]) were changed to weaker hexamers (GTCCAA [Rank 241, SSE-0.038] / TGCGAG [Rank 217, SSE-0.037]; Supplementary Table S13-S14) and transfected the constructs into HEK293T cells. We analyzed mCHERRY fluorescence and performed RT-PCR (Fig. 5B). Changing the hexamers abolished splicing at these splice-sites but revealed cryptic 5’ and 3’ splice-sites that resulted in an inclusion of 26bp (additional 8 bp from the alternative donor site and 18 bp from the alternative acceptor site), which caused frameshift disrupting mCHERRY fluorescence.

Second, as a weak intron we selected intron 2 of *CALM3*, (chr19:46,608,327-46,608,481) and changed the donor hexamer from GTGGAT [Rank 200, SSE-0.117] to GTGAGT [Rank 3, SSE-0.724]. With the original CALM3 intron, there is an alternative donor 13bp downstream (GTGAGC [Rank 9, SSE-0.653]) is used in splicing, which disrupts mCHERRY expression due to a frameshift (Fig. 5C). However, change of the hexamer resulted in proper splicing with mCHERRY fluorescence (Fig. 5C).

Third, we selected *MYO15B* intron (chr17:75592302-75592463) where two acceptor sites at chr17:75592427 and chr17:75592463 compete while the donor is less variable. GWAS analysis showed that the splice-site choice mapped to an SNP at chr17:75592461 (A/G) in the hexamer sequence (TACAAG [Rank-148, SSE-0.279] / TACGAG [Rank 201, SSE-0.049], Supplementary Table S5). We changed the sequences at these competing sites to have various combinations of hexamers with differing strengths (Supplementary Table S13-14). Confirming the causality in our GWAS results, a change of A to G reduced the use of chr17:75592463 and promoted the use of chr17:75592427. Analysis of diverse combinations revealed that the highest mCHERRY expression and proper splicing was observed with the construct where we supplied the donor and acceptor sites with higher ranked hexamer sequences (Supplementary Figure S16A, B). In all other cases we observed sub-optimal splicing, including the use of alternative splice-sites (Supplementary Figure S16A, B).

Finally, we took our synthetic worst intron and modified it to assess minimal features that could improve splicing. Adding a polypyrimidine tract and a branch point with consensus sequences had negligible effects suggestive of additional requirements (Supplementary Figure S17). When we also changed both donor and acceptor hexamers into stronger ones, we could observe some level of proper splicing (Supplementary Figure S17). Taken together, these findings experimentally demonstrated that the hexamer sequences are among the primary determinants of splice-site choice.

## Discussion

We have demonstrated that there is extensive, genetically determined variation in splicing in diverse organisms, which can be precisely mapped to the specific, often causal, nucleotide variation by combining splice-site strength with GWAS. Using the individual splice-site strength as a phenotype for GWAS allowed us to specifically link the impacts of genetic variation to the usage of specific donor or acceptor splice-sites. Often, a mutation that directly impacts the strength of one splice site will have indirect effects on the usage of other sites. For example, in case of the *MYO15B* (Supplementary Figure 16A, B) an SNP at chr17:75592461 reduces the efficiency of the splice acceptor site 2bp downstream of the SNP (chr17:75592463), but also indirectly increases the efficiency of the competing acceptor site at chr17:75592427. We were able to not only demonstrate that we could map splice-site usage of both competing sites to allelic variation at chr17:75596461 (Supplementary Table S5) but also confirm this through minigene assays (Supplementary Figure S16A, B). Thus, we can differentiate direct and indirect effects in many instances, simply from the distance of the associated SNP from the splice-site. Differentiating such primary and secondary effects is harder to achieve with other approaches. Thus, we would argue that our approach focused on individual splice-sites would provide a better understanding of the regulation of splicing as opposed to other approaches based on isoforms or splicing events.

We have shown that common splicing variation is primarily driven by “*cis”* rather than “*trans”* regulatory genetic variation. While it is clear that there are several trans-acting factors that play a critical role in splicing (Lee and Rio, 2015), natural variation in “*trans”* effects (i.e., a change in splicing driven by a trans acting factor) could be brought about in two different ways (Ule and Blencowe, 2019): there can be sequence variation in a trans-acting factor modifying its function, or there can be a change in the potential binding site of a trans-acting factor. Both instances will result in splicing variation. While the former is likely to have an impact on multiple genes, the later allows specific changes restricted to individual genes/splice-sites, generating variability which can be beneficial in evolution (Singh and Ahi, 2022). Our findings support the idea that *cis* regulatory changes are preferred over *trans* regulatory sequence variation. We, however, acknowledge that due to our highly stringent analysis (i.e., having higher thresholds used peak calling and primarily looking at the top or closest associated SNP only), we could potentially underestimate the impact of *trans* variation, particularly if the effect size is relatively small. Confirming the causality of these *trans* associations would require further experiments.

We have presented a universal core logic that explains splice-site choice in eukaryotes based on hexamer rankings. Our findings that (1). Hexamer prevalence at splice-sites is correlated with splice-site strength rather than genomic prevalence, (2). Hexamer ranks explain splice-site choices, and (3). Hexamer ranks correlate across diverse species together argue for evolutionary selection for hexamer sequences.

Perhaps unsurprisingly, but now clearly demonstrated, we have shown a near perfect correlation with sequence distances from U1 snRNA base-pairing site to the average strength of intronic hexamers at splice donor sites (Supplementary Figure S15B); the same intronic hexamers whose ranking explains much of the splice site choice at a population level. The donor site is typically bound by U1 snRNA and sequence variability could potentially lead to differential binding kinetics of U1 snRNA and its associated proteins, with potential consequences to splice-site choice (Rogalska et al., 2023). In addition to snRNA-RNA base-pairing some of effects could also be attributed to hexamers (e.g., GTCTTT, GTCTTA) being binding sites for proteins such as PTBP1 (Giudice et al., 2016), which may interfere splicing when the sequences are at the 5’ donor sites (Sharma et al., 2011). Consistent with this, we found the average strengths of these hexamers to be very low (Supplementary Table S13). On the other hand, we observed hexamer GTAACG, which is a known binding motif for DAZAP1 (Giudice et al., 2016), a protein known to promote splicing, to be with a high average strength. Thus, in addition to snRNA-RNA base pairing, RNA binding proteins could influence the average strengths. At the acceptor sites, where RNA-protein interactions play a critical role in splice-site selection (Mendel et al., 2021; Rogalska et al., 2023; Smith et al., 1993), we observed similar correlations between the sequence distances from the highest ranked hexamer and the average strengths (Supplementary Figure S15C). This can reflect differential binding of proteins with RNA. These differential interactions, while requiring further validation, could be the potential underlying mechanisms through which hexamer sequences influence splice-site choices.

Typical GWAS analysis results in the identification of genomic regions associated with phenotypic variation. Over the past 15 years, with the explosion of GWAS studies, there are many genetic variants of interest that have been catalogued to be of significance (Abdellaoui et al., 2023). However, understanding the implications of these SNPs and the underlying mechanisms by which they act remains a huge challenge (Lappalainen and MacArthur, 2021). Even with the highest associated SNPs from SpliSER-GWAS, more than 10% have been identified to be variants of interest for diverse phenotypes (Supplementary Table S16). This suggests splicing as a potential molecular mechanism that could be explored in the context of these phenotypes, with important implications for personalized medicine.

In summary, we have presented a core principle that explains splice-site choice across eukaryotic organisms. We have shown that the core determinants of splicing across diverse species is intronic hexamer sequences, and that these are the major source of splicing variability at a population level. Our catalogue of associations in Arabidopsis, Drosophila and Humans can be further exploited both for functional analysis of pathways of growth, development, or environmental responses. The hexamer rankings that we have presented can be used for engineering specific splicing outcomes through gene editing techniques for desirable phenotypes of agricultural and/or medical relevance across eukaryotes.

## Acknowledgements

We thank the GTEx Consortium, 1001 Genomes Project and the Drosophila Genotype Reference Panel project and the researchers who made the transcriptome datasets available for scientific community use, which allowed us to undertake a cross-species approach. We thank Elaine Zhang, Hongyu Zheng, Shuyu Yan, Dhruv Sheth, Tenghao Zheng and Matthew Parker for their help and discussions and the members of the SKB/GATC labs for their critical comments on the manuscript. CID, JG, and JC are supported by an Australian Government Research Training Program. MG is supported by an Indonesian Government Education Scholarship (Ref No. 202101030416). JR is supported by a Victoria Cancer Agency fellowship (Grant Number: MCRF20035). This work is supported by Australian Research Council Discovery Project DP190101479 (SB), Australian Research Council Future Fellowship (FT190101818) (SS), National Health and Medical Research Council Ideas Grant APP1182090 (SB & SS).

## Author contributions

Conceptualization: SB; Methodology: CID, SP, AB, JG, AC, JR, MDA, PPD, AFL, SS, DP and SB; Data curation: CID, SP, AB, JG, AC and SB; Software: CID, SP, AB, AC, AFL, DP and SB; Formal Analysis: CID, SP, AB, JG, AC, SM, JC, MG, RDS, RB, SS and SB; Investigation: CID, JG, JC, MG, RDS, RB, SS and SB; Writing – Original Draft: CID, SP, AB and SB; Writing-Review and Editing: CID, MG, SP, AFL, YLG, SS, MDA, RB and SB; Visualization: CID, SP, AB, AC, JC and SB; Supervision: JR, YLG, AFL, RB, SS, DP and SB; Project Administration: SB; Funding acquisition: SS and SB.

## Materials and Methods

### DNA/RNA analyses

DNA and RNA extractions from plants, flies and HEK293T cells were carried out as described previously (Eimer et al., 2018; Zhang et al., 2021). Plants were grown as described previously (Eimer et al., 2018). DGRP panel of flies were gifted by Prof Charles Rubin (University of Melbourne) and RNA was extracted as described previously(Zhang et al., 2021). Fly stocks were maintained in standard fly media until DNA/RNA extractions. HEK293T cells were cultured as per standard procedures in DMEM media with 10% FBS and 2mM glutamine. To analyze splicing, 1μg total RNA was converted into cDNA and the splicing patterns were analyzed using primers listed in Supplementary Table S17. All standard molecular biology works were done as described previously (Eimer et al., 2018). All constructs were sequence verified before their downstream uses. RT-PCR products were gel purified and sequenced to confirm specific splice junctions.

### RNA-seq data alignment for processing

The Arabidopsis 1001 genomes RNA-seq data was aligned with TopHat2 and splice junctions were identified with Regtools as previously described (Dent et al., 2021). Briefly, 6854 RNA-seq samples were downloaded representing 728 accessions (PRJNA319904) (Kawakatsu et al., 2016). Reads were aligned to the TAIR10 genome using TopHat2 (v2.1.1; paramete– -- minIntronLength 20, --maxIntronLength 6000, -p 6) (Kim et al., 2013). For Drosophila, transcriptomic data was obtained from the Drosophila Genetics Reference Panel – DGRP2 (Huang et al., 2015). The 957 DGRP2 RNA-seq samples were downloaded, and the data was aligned using STAR version 2.7 (Dobin et al., 2013) to the BDGP6 reference genome (paramete– --outFilterMultimapNmax– --alignSJoverhangMin– --alignIntronMin 20 –alignIntronMax 150– -- outSAMtype BAM SortedByCoordinate). Aligned bam files were indexed with Samtools version 1.12. Regtools was used to generate gap junction files (parameters-regtools junctions extract -a 6 -m 20 -M 15000 -s 0). For the human data 372 BAM files representing human heart atrial tissue generated by the GTEx project (Consortium, 2013) were downloaded. A BED file also known as a gap junction file, containing a catalogue of splice junctions detected in the alignment was generated with Regtools junction extract (parameters -m 20 -M 16000 -a 6 -s 0 -o ${SAMPLE}.bed {SAMPLE}.bam).

### Quantification of splice-site strength/usage estimates (SSE)

The resulting BAM and splice junction BED files were processed with SpliSER v0.1.8(Dent et al., 2021). We filtered sites which had at least 10 reads crossing the splice-site in at least three replicates, in at least 100 accessions and taken them for further analysis (Table 1). Fly data was split into female SSE (fSSE) and male SSE (mSSE) datasets. For each splice-site, SSE was averaged across samples of the same genotype and sex if at least two replicates passed the previous filtering steps.

Variability in splice-site usage between individuals was evaluated in two different ways. First, the variance of the distribution was calculated and individuals who fell in the upper quartile were taken for further consideration. Second, we computed the range in splice-site usage among the individuals in each of the species. Typically, there was a correlation between the two measures. Broad-sense heritability (H^2^) was calculated from the averaged SSE phenotypic values as the proportion of total variance attributed to the variance between the DGRP/1001 project accessions, using one-way ANOVA with the genotypes as a factor and SSE as a response. Splice-sites were ranked by their H^2^ and total variance in SSE, and sites in the top quartile for both H2 and total variance were taken for GWAS mapping. For the human heart data, all sites that were in the upper quartile of variance were taken for analysis since heritability could not be calculated due to the absence of replicates.

### GWAS and peak-calling from Manhattan plots

We ran GWAS for each site using GEMMA v0.98.3 (Zhou and Stephens, 2012), using genotypic data from 1001 Genome project/DGRP project or the GTEx project. A distance square matrix kinship file was generated using the VCFs and then utilized in GWAS analysis. Only the variants that had a minor allele frequency greater than 5% were considered for the association study.

Initially, we manually called the peaks for Arabidopsis and then automated the process through benchmarking with Manhattan Harvester (Haller et al., 2019), with a minimum peak SNP count of 50. To account for the global noise in Manhattan plots in this approach, we additionally required that the –log10(pValue) of the top SNP of a peak must lie 1.33 times higher than the average of the top 5 SNPs in the plot (with none of the 5 being within 750 kb of each other) for the peak to be included. From each of the called peaks, a single SNP with the most significant p-value was selected as the top SNP. In cases where multiple SNPs had the maximal association significance, the one closest to the splice-site was chosen as the top SNP and selected for downstream analyses. All selected associations of a SNP and a splice-site were compiled into a comprehensive SNP table (Supplementary Tables S3 – S5). The distance of each top SNP from the associated splice-site was calculated and normalised to the splice-site position and strand, with position zero being the first base in the intron for the donor site (“G” of “GT”), and last base of the intron for acceptor sites (“G” of “AG”). Associations were characterised as “cis” if the top SNP was within 1Mb from the associated splice-site; all other associations were labelled as “trans”. Allelic change in SSE (ΔSSE) was calculated as the average major allele SSE subtracted from the average minor allele SSE. Percentage Variance Explained (PVE) of SSE by each SNP was calculated as per Shim et al (Shim et al., 2015).

### Single nucleotide effects on splicing variation

We calculated the proportion of variance attributed to each position around the splice-site as the ratio between within-group variance and between group variance using this formula.

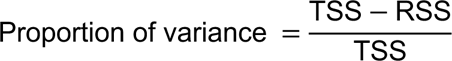

where TSS is the sum of squares of Splice-site Strength Estimates (SSEs) across all splice-sites, and RSS is the sum of the sum of squares of SSEs of sites with each nucleotide at the given position. We calculated this for all splice-sites in Arabidopsis, Drosophila and Humans, taking the information from multiple individuals that were used in the GWAS analysis.

### Second-order effects of nucleotides around splice-sites

For each species in the GWAS dataset we sampled down to approx. 2 million splice site/sequence combinations. We investigated the sequences -7 upstream until +3 downstream of Acceptor Sites, and -3 upstream until +7 downstream of Donor sites. Preliminary analysis with two-way-ANOVA suggested that all positions around the splice site were significantly associated with changes in splice site usage, and that there were significant interaction effects of all nucleotides at all positions. To capture second-order effects (pairwise interactions of nucleotides which differ from a naive additive model) we calculated the mean splice site usage of each splice site with either nucleotide 1 (e.g., Adenine at position -7) and nucleotide 2 (e.g., Guanine at position -6; together forming an AG upstream of the splice site). We subtracted the mean usage of sites with each nucleotide from the grand mean, and added the differences together to form an naive expectation of their interaction if it were additive (e.g., -7A = -0.1, -6G = -0.15, so we expect sites with -7/-6 AG to have an average usage 0.25 below the mean; capped such that splice site usage could not exceed 0 or 1). We then looked at the difference between this expectation and our actual observations. We plotted all these differences in sorted order and identified outliers as those sitting beyond the elbows of the resulting plot (Fig 3E-F, Supplementary Figures S10). Elbow points were calculated as the points furthest from the lines drawn between the mid-point and either extreme of the plot, to which we added a further conservative buffer of 0.1 (or 10% change in splice site usage beyond the elbow threshold). The pairwise interactions values are given in Supplementary Tables S10-S11 and the heat maps of interactions for each species is shown in Supplementary Figures S11-S13.

### *k*-mer effects on splice-site choice

To identify an ideal *k*-mer length, we assessed the ability of the *k*-mer to explain splice-site choice in the following manner. We scanned +/-100 or 200bp from the splice-site for all GTs for donors and AGs for acceptors. We then extracted the *k*-mers surrounding the GT/AG in the window taking into account of the existing mutations using corresponding VCF files for Arabidopsis, Drosophila and Humans and compared their average splice-site strength to generate *k*-mer ranks. The number of occasions when the splice-site was having the strongest and unique *k*-mer among the *k*-mer for all GT or AG in that window were noted. This number over the total analyzed sites to obtain a success rate. We then adjusted this score based on the percentage of possible *k*-mers that are present in splice-sites to obtain the percentage of splice-site choices explained the *k*-mer (Fig 4A) for all three species. We excluded splice-sites that are within 100/200 bp of each other, eliminating the competition between detected sites for this analysis.

### Identifying the best nucleotides for each position that promote/suppress splicing

From the SNP table, we computed the distances of the highest and closest associated SNPs from their corresponding splice-sites. The table was filtered to only include splice-sites for which the highest associated SNP fell within 100bp from the splice-site. We then trimmed down the list so that each associated SNP was giving information only for a single site by taking the closest splice-site. This resulted in a unique set of associated SNPs, each of which gave information about one unique splice-site. Subsequently the list was separated into donor and acceptor sites and the data was processed to produce the distribution graphs for each species. The data from all three species was combined and for each association the splice-promoting nucleotide and the splice-reducing nucleotides were deciphered as the most frequent nucleotide seen for greater or lesser usage of the splice-site. To synthesize the best and worst introns, we carried out this analysis on acceptors from -50 to 0 and from 0 to 50 for donors and stick them together as shown in Supplementary Table S6-S7. We included GT and AG and a branch point adenosine in all constructs in this analysis. The worst construct was then modified to include either a polypyrimidine tract (PPT) or a PPT and a consensus branch point or a PPT, a consensus branch point, and strong hexamers to assess the importance of hexamers. For all splice-sites across the genome, we computed the average SSE for each of the 4 possible nucleotides at each position relative to the splice-sites and plotted the means as a violin plot and considered the nucleotide with the highest mean as the best one to promote splicing for that position.

### mCHERRY reporter construct analysis

To assess the splicing impacts experimentally, we interrupted the mCHERRY ORF of pGH044 (Addgene Plasmid #85412, RRID: Addgene_85412). We scanned mCHERRY for “AGGT” stretch and placed the intron between the AG and GT. We synthesized the intron with a part of mCHERRY through commercial suppliers (IDT-Australia) and then used restriction cloning to replace this cassette containing the intron within the plasmid pGH044. All constructs generated are listed in Supplementary Table S7. Sequence-verified constructs were transfected into HEK293T cells using Lipofectamine^TM^ 3000 (Invitrogen). Cells were visualized under a fluorescence microscope. RNA from transfected cells were extracted for RT-PCR analysis.

### Hexamer analysis on diverse species

We downloaded the raw RNA-seq data (.fastq) of different species from SRA using SRAtoolkit. The RNA-seq reads were aligned to their corresponding genome annotations from SRA using STAR. Subsequent .bam files were then indexed and regtools was used to generate .bed files, detailing the splice junction information. Then, SpliSER was used to calculate splice-site usage. SpliSER’s output file was used as input for a custom-made python script that identified hexamers for each splice-site and calculated the average Splice-site Strength Estimate (SSE) for each hexamer. For Arabidopsis, Drosophila and human data, the entire dataset containing all individual samples (e.g., 200 genotypes in Drosophila) were used to generate obtain the hexamer for all sites, which further aided in obtaining a robust estimation of the SSE or count. For all other samples, single RNA-seq data was used to obtain the SSE or count ranks. These ranks were then used to assess the percentage of splice-site choices explained by the hexamers in each of these species. All the data from diverse species is reported in Supplementary Table S12-S14. This data was used to obtain pairwise correlations between species through custom-made python scripts. The results are reported in Supplementary Table S15. To analyze species wide correlations, we computed the *p-distance* using MEGA software (Tamura et al., 2021). After grouping hexamers by *p-distance*, we computed the average SSE for each group by averaging the SSE of all hexamers within the group.

### Analysis of the Rosenberg et al. minigene constructs data

We used the RNA-seq data generated by Rosenberg et al (Rosenberg et al., 2015), which was downloaded from NCBI in FASTQ format. As the reads were multiplexed, we used fastq-multx (https://github.com/brwnj/fastq-multx) to allocate sequences to different minigene constructs based on the barcodes. The de-multiplexed reads were aligned to the corresponding reference sequence using STAR. These aligned reads were processed with SpliSER (v1.8) and SSE values were obtained for all the detected splice-sites from all 265,137 constructs. Custom-made Python scripts were used in downstream analysis, which involved extracting the hexamer sequence information from the reference sequence and identifying all possible competing pairs within each construct. Subsequently, we calculated the average SSE value for all hexamers in each competing pairs and determined the proportion of winning hexamer based on this average SSE value.

## Supplemental information

**Supplementary Figure S1.**
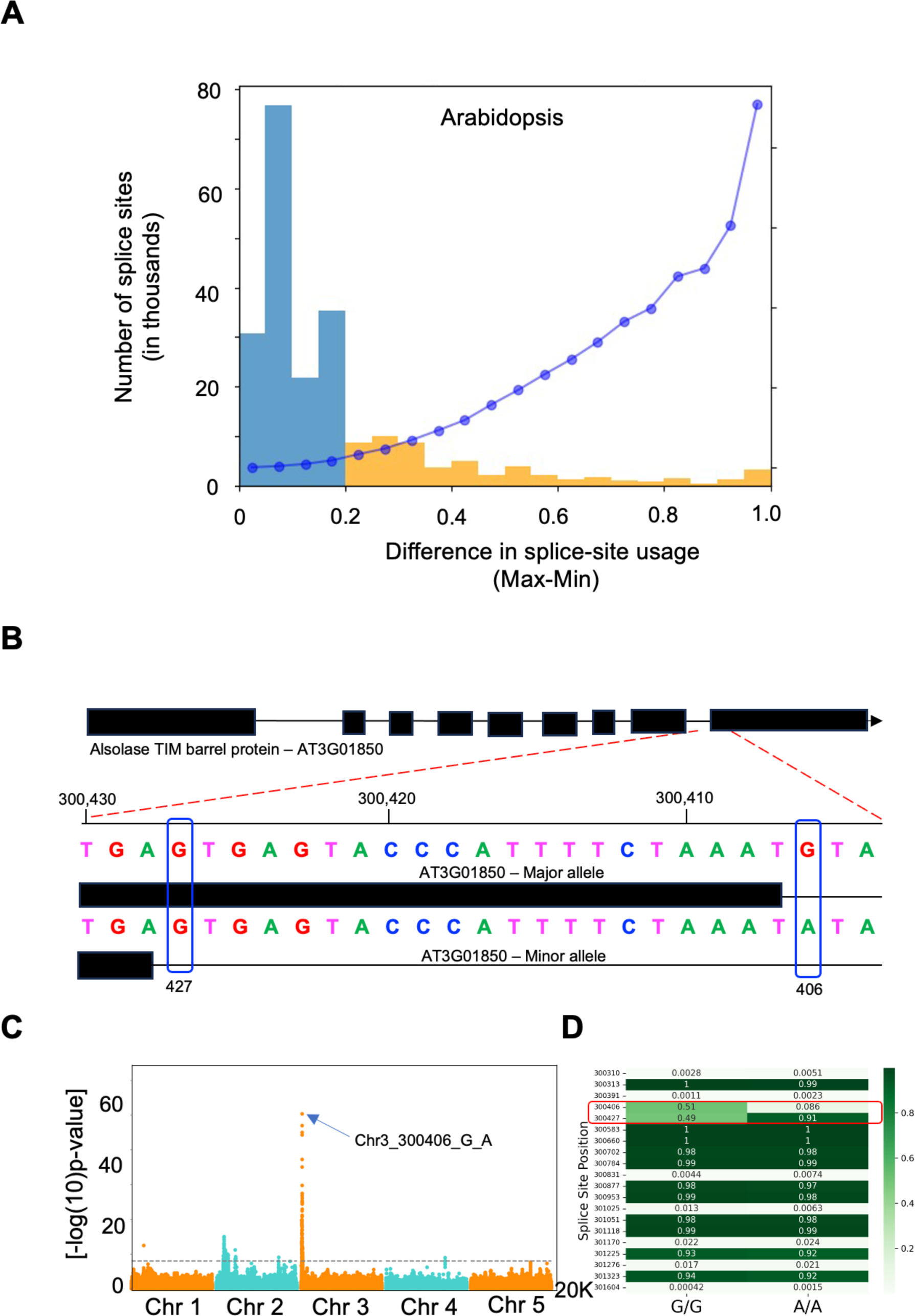
There is extensive variation in splice-site usage, which can be mapped accurately with SpliSER-GWAS. A) Distribution of splice-sites that display variation between individuals in Arabidopsis. Yellow region highlights the sites that display more than 20% variability between extreme samples. Blue dots represent the average variance for each of the bins. B-C) SpliSER-GWAS identifies the causal SNP for variation in the usage of splice-sites at the AT3G01850 locus in Arabidopsis. The splice-site mutation at 406 allows the usage of 427 as a splice-site and variation in the usage of 406 as well as 427 maps to the 406 polymorphism. D) The effect of the mutation is specific to this site and does not affect neighboring sites.

**Supplementary Figure S2.**
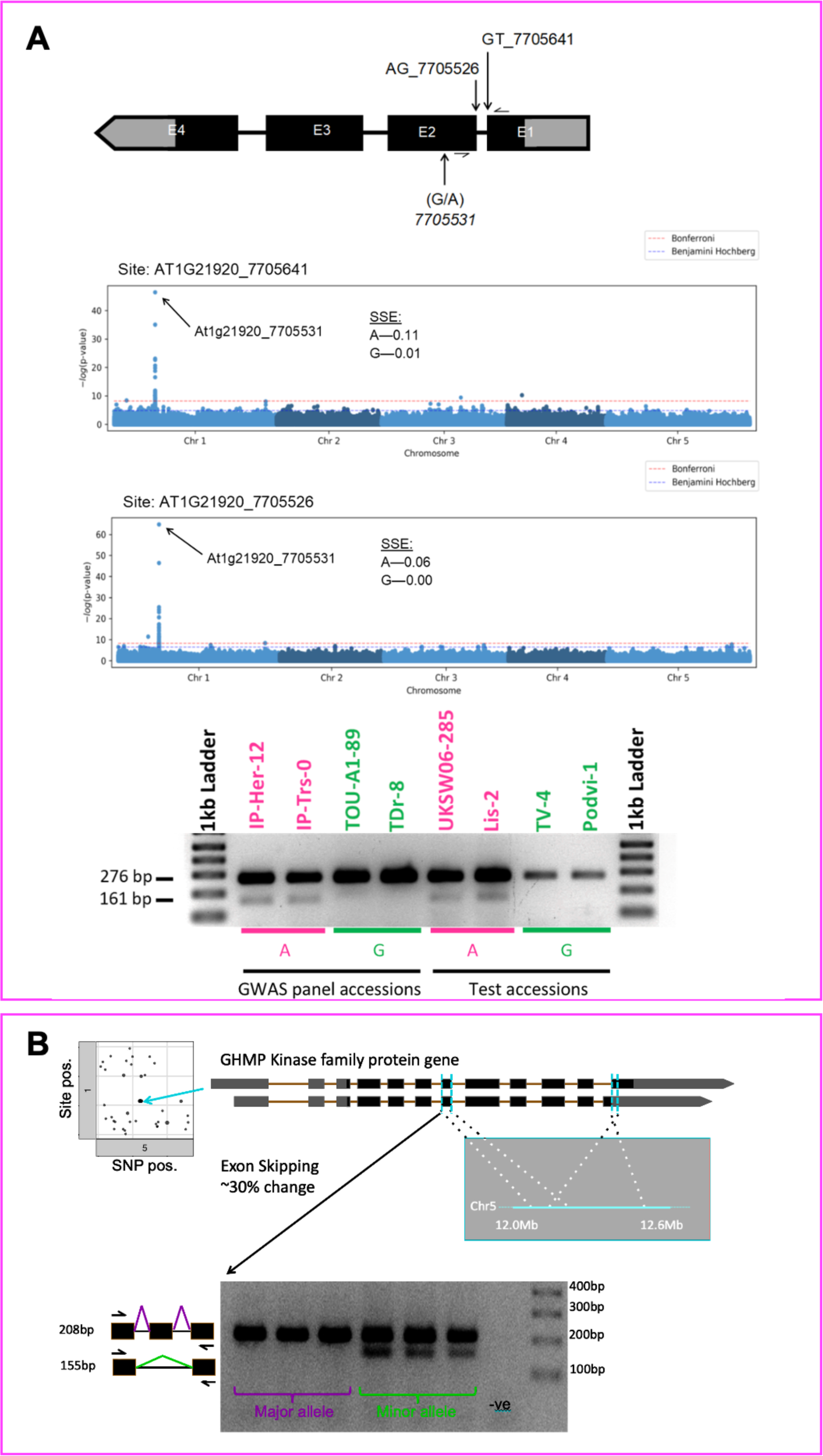
SpliSER-GWAS associations are experimentally verifiable. A) Mapping of a minor-effect intron splicing at AT1G21920, which encodes a histone methyl transferase and natural variation of an SNP at position 7705531 is associated with the use of both splice-donor and acceptor sites. The presence of “G” at this position abolishes splicing (larger bands) and presence of “A” leads to partial splicing of the intron resulting in two bands. Test accessions that are not part of the GWAS panel display the same splicing patterns confirming GWAS associations. B) Experimental verification of a “*trans”* association in *Arabidopsis thaliana.* An exon-skipping outcome that occurs due to differential usage of few different sites in a gene in Chromosome 1 that encodes a GHMP kinase family protein maps to a region in Chromosome 5. Several “*trans”* associations map to the same region. RT-PCR analysis with specific genotypes confirmed the association, though the mechanisms remain unclear.

**Supplementary Figure S3.**
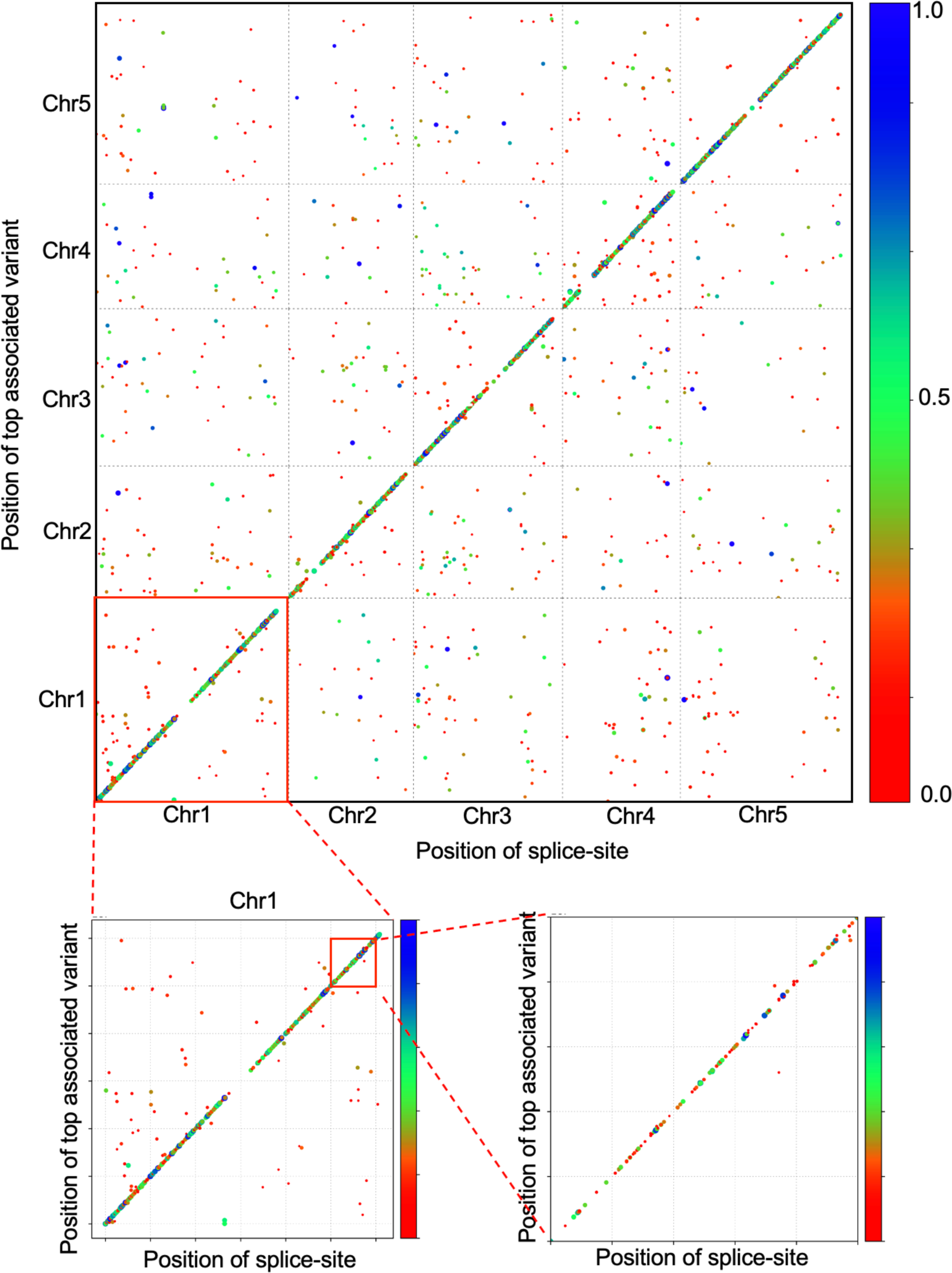
Splicing variability is mostly “cis” regulated in Arabidopsis. A-C) Scatter plot of splice-site positions and their highest associated SNPs in the Arabidopsis genome. Chromosome 1 is zoomed in at two different levels to show the *cis* nature of associations. Colour scale represent the percentage of variance (PVE) explained by the associated top SNP. The sizes of the dots are also correlated with PVE.

**Supplementary Figure S4.**
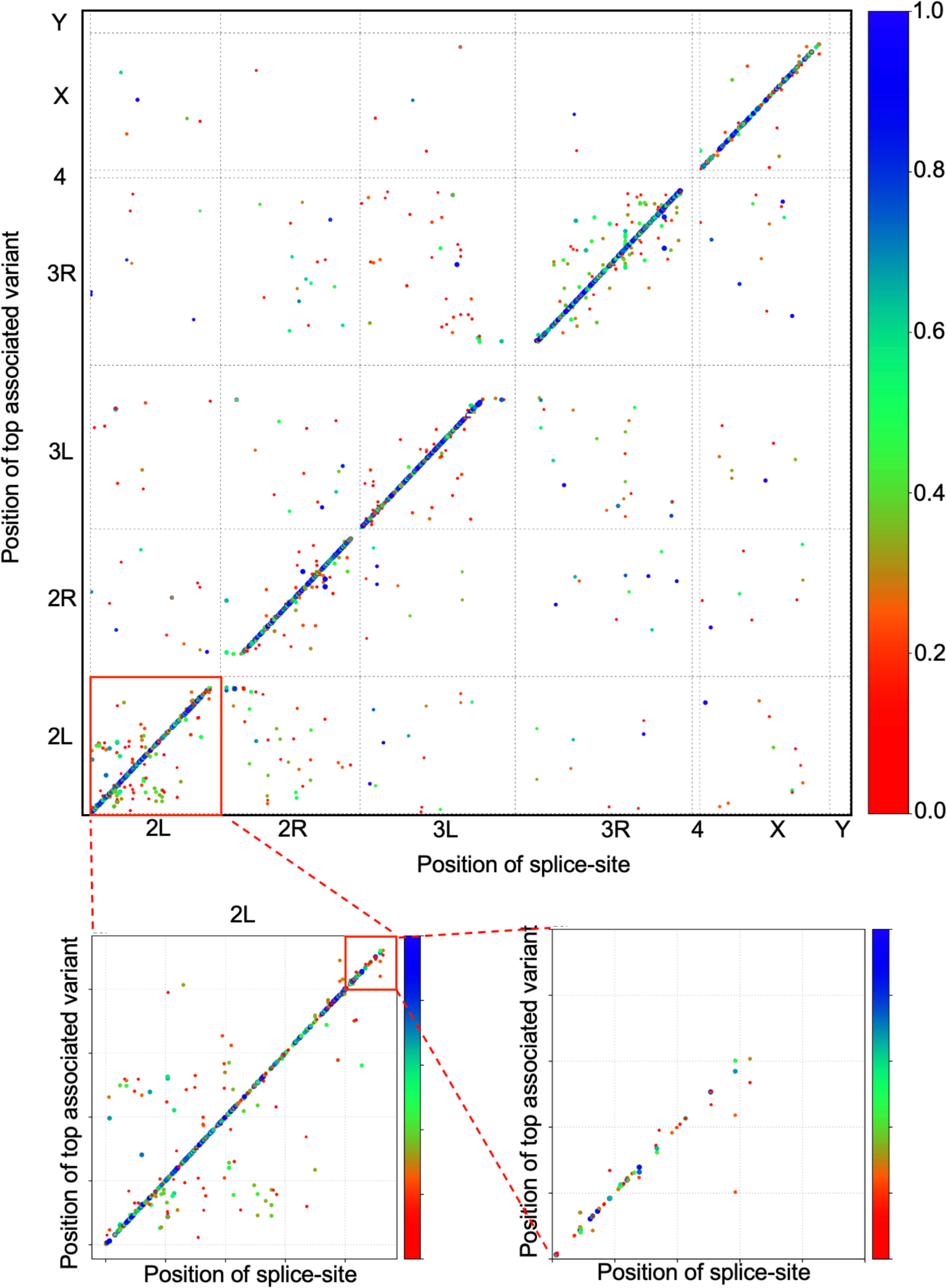
Splicing variability is mostly “*cis”* regulated in Drosophila. A-C) Scatter plot of splice-site positions and their highest associated SNPs in the Drosophila genome. Chromosome 2L is zoomed in at two different levels to show the *cis* nature of associations. Colour scale represent the percentage of variance (PVE) explained by the associated top SNP. The sizes of the dots are also correlated with PVE.

**Supplementary Figure S5.**
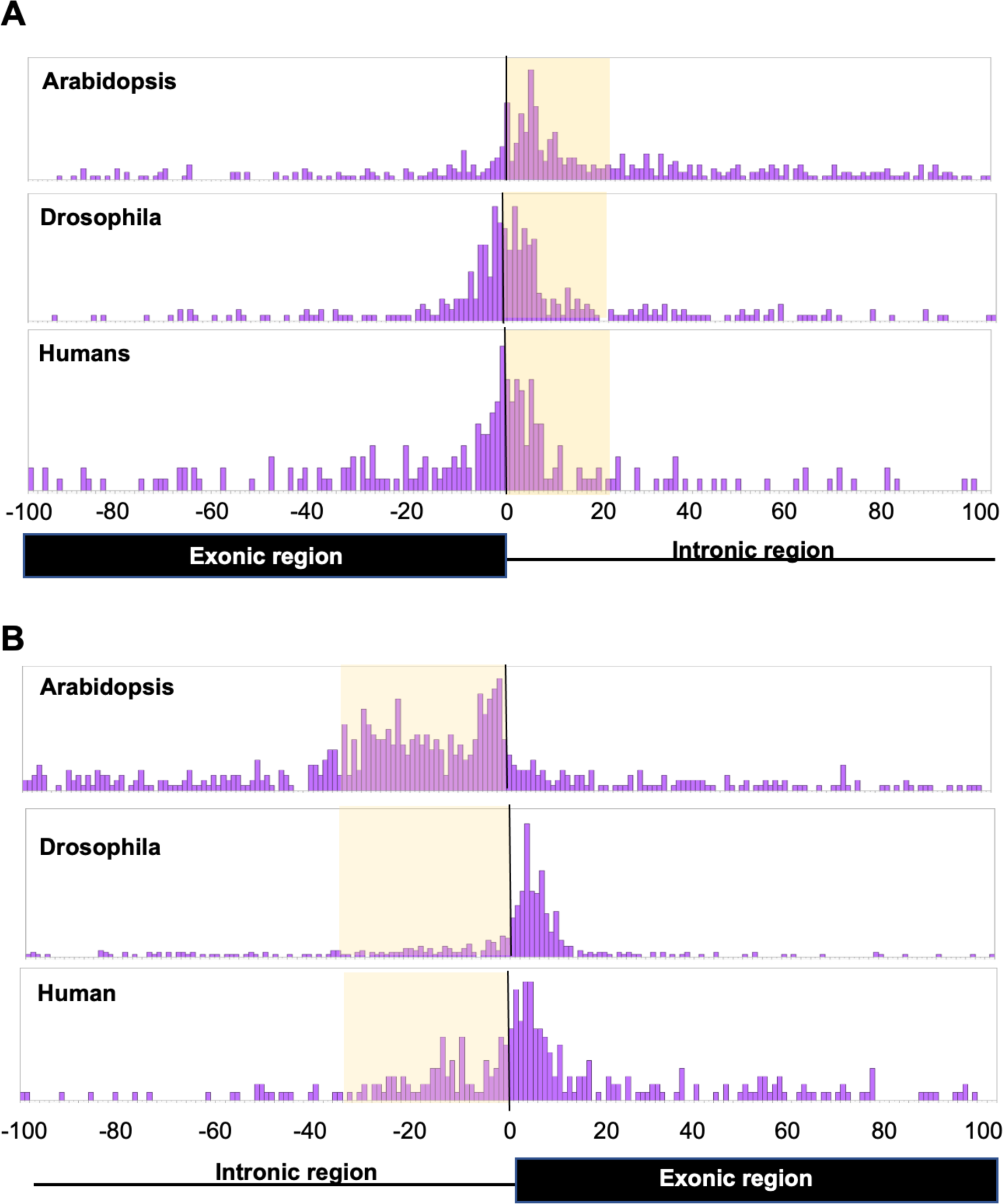
Genetic variation affecting splice-site choice is often near the splice-site. A-B) Distribution of the distances of the closest associated SNPs detected in SpliSER-GWAS for donors (A) and acceptors (B) in Arabidopsis, Drosophila and Humans. The “G” of “GT” or “AG” is plotted as position 0. Intronic regions with higher number of associations is shaded for clarity.

**Supplementary Figure S6.**
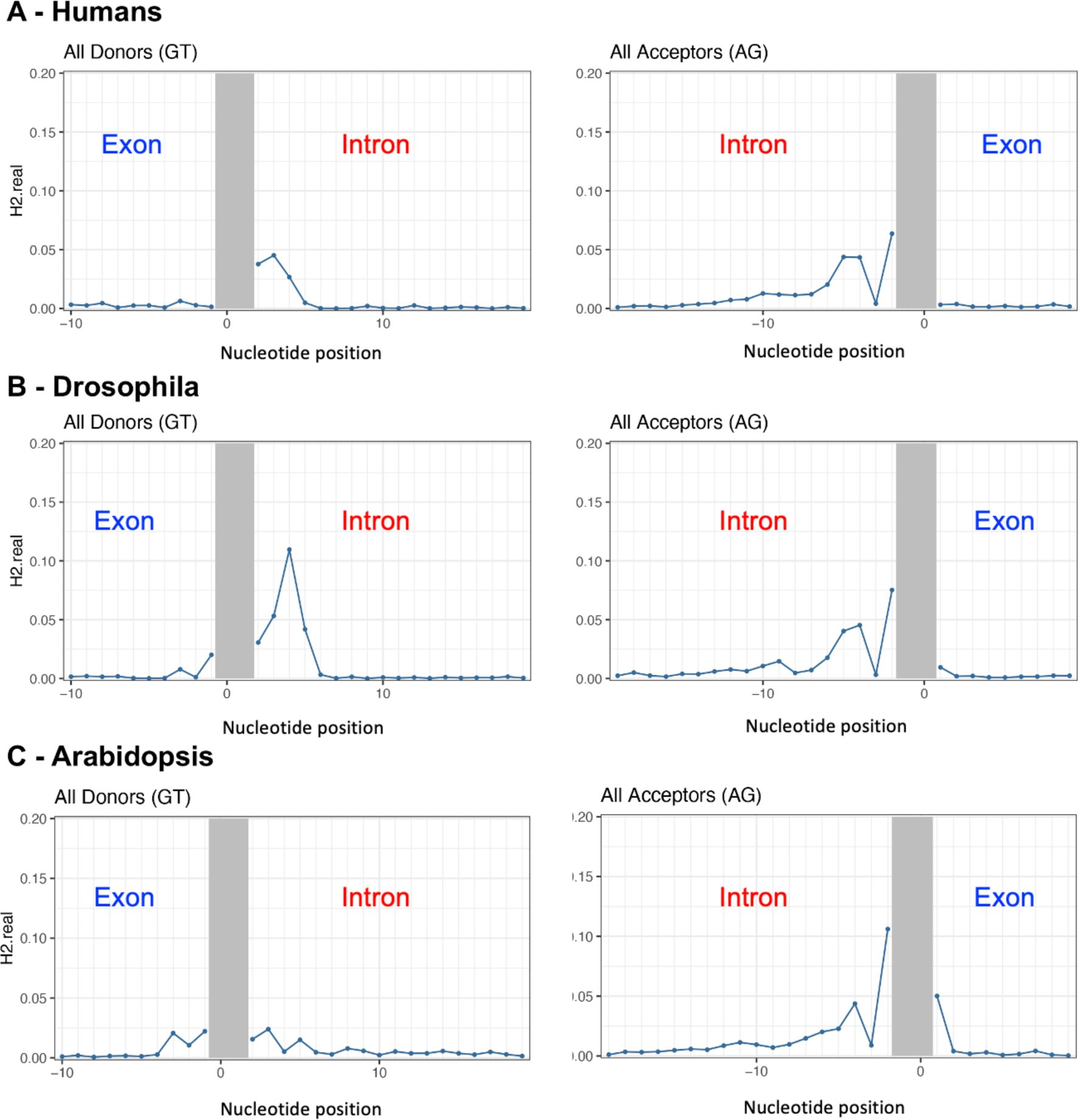
Intronic variability is the primary driver of variability in splice-site strength. Realized heritability for every single nucleotide position surrounding splice-sites in Humans (A), Drosophila (B) and Arabidopsis (C). The grey region represents GT or AG, which do not have any polymorphisms, since the analysis is restricted to the GT or AG sites. The intronic regions in all three species explain more variability than the exonic regions.

**Supplementary Figure S7.**
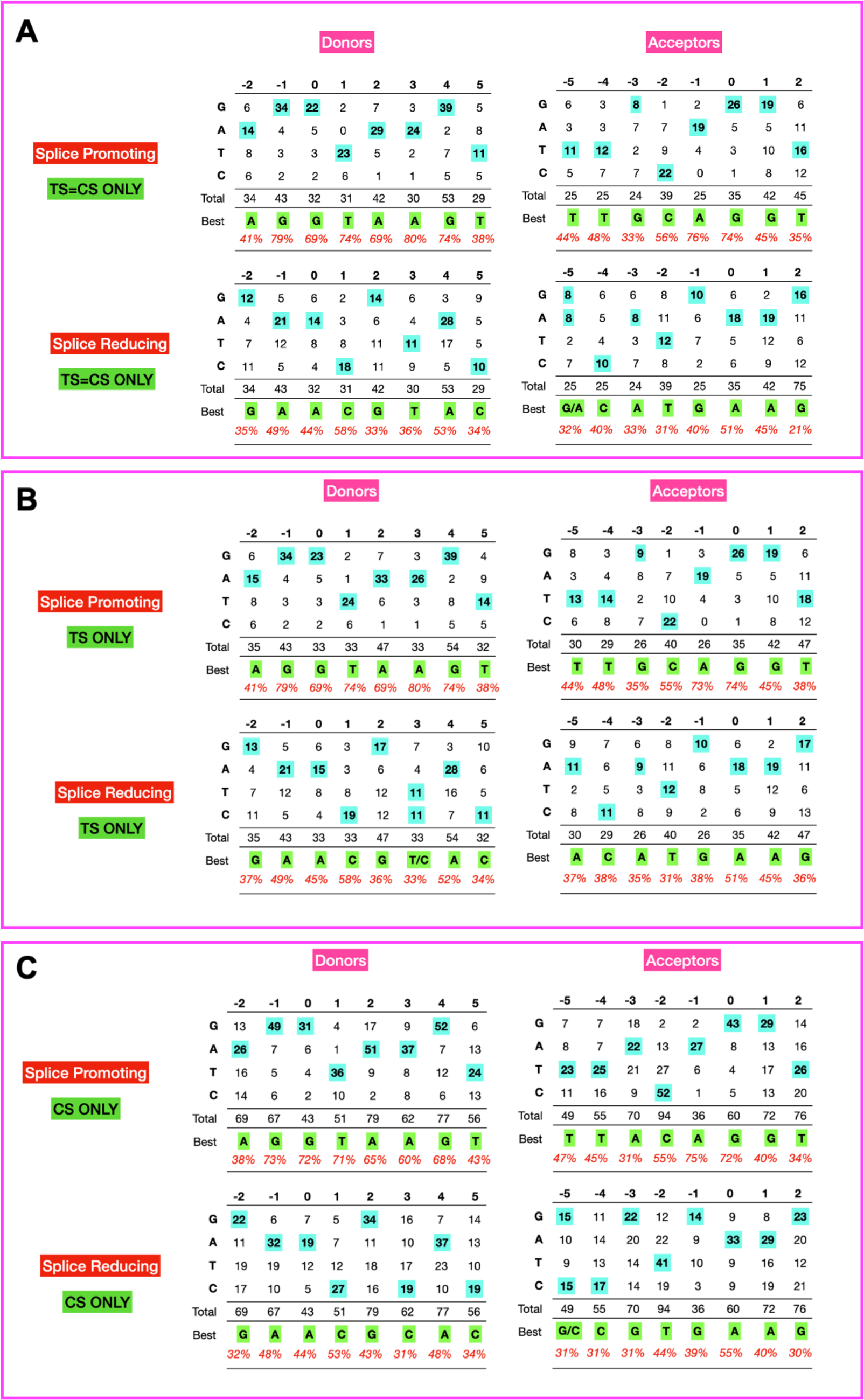
High resolution SpliSER-GWAS allows inferring best nucleotides that promote splicing. The distribution of splice-promoting allelic variation for positions -2 to +5 around the splice-donor site and -5 to +2 around the splice-acceptor site based on associations from all three species considering either Top SNP (TS) (B), the closest SNP in the peak (CS) (C), or where both the CS and TS are the same (A). The most frequent splice-promoting nucleotide is highlighted for each position. For control to account for general sequence variation, splice-reducing nucleotides are also shown.

**Supplementary Figure S8.**
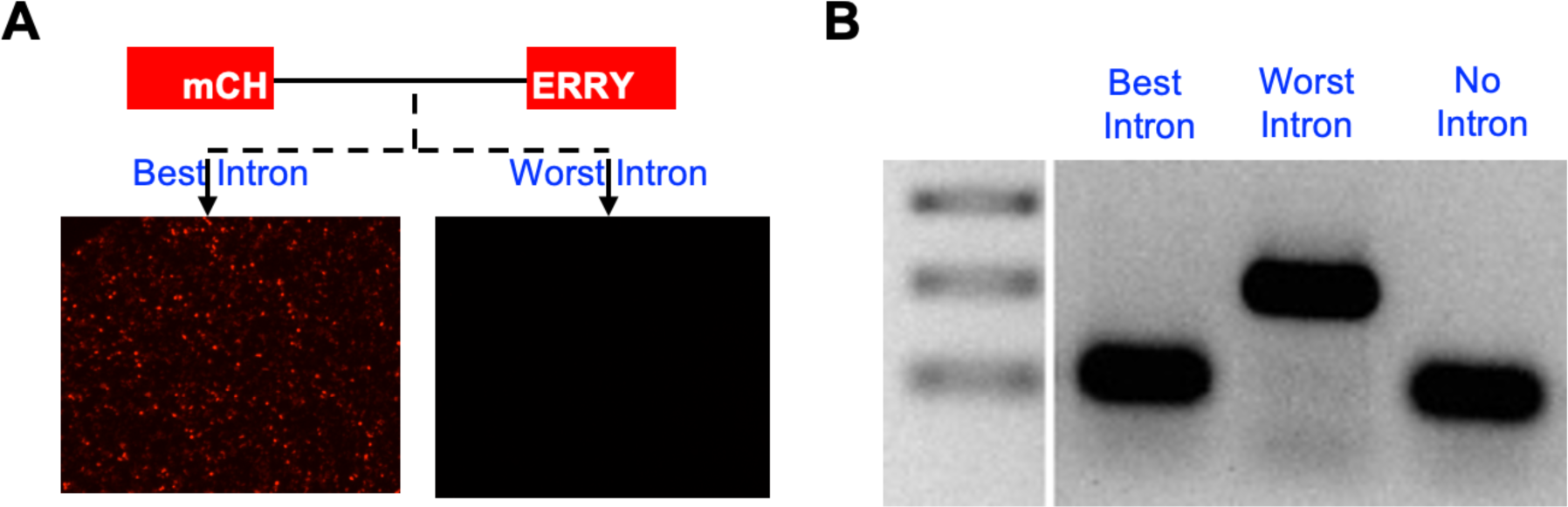
Experimental verification of the performance of synthetic introns designed from the best and worst nucleotide combinations through an mCHERRY mini gene assay. Effect was assayed through mCHERRY fluorescence (A) and via RT-PCR (B).

**Supplementary Figure S9.**
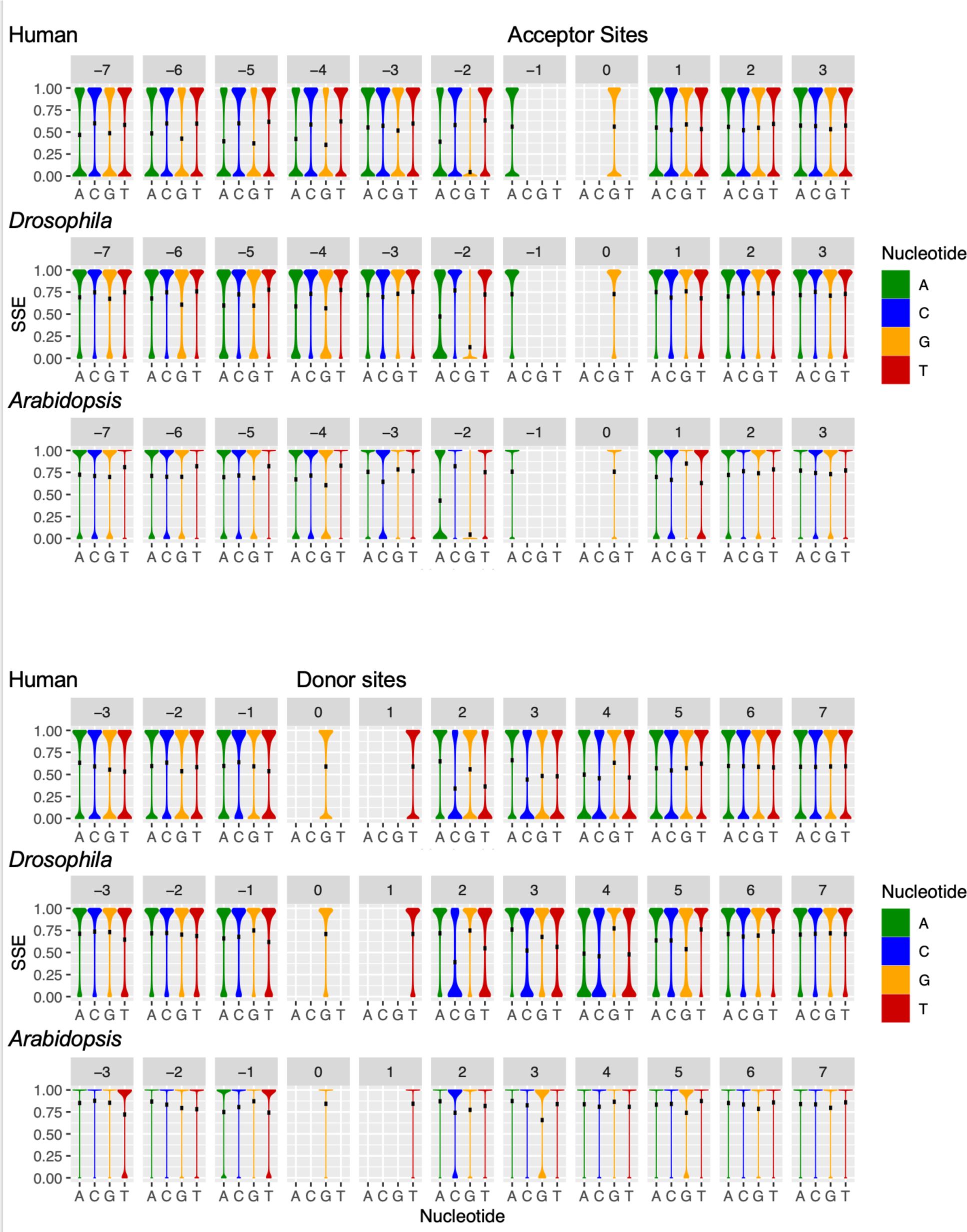
The mean Splice-site Strength Estimate (SSE) of splice sites harbouring each of the four possible nucleotides at each position around the splice site. Top – Splice acceptor sites (AG only) from position -7 to +3. Bottom – Splice donor sites (GT only) from position -3 to +7. Black dots represent the mean. Downsampled in each species to 1-2 million site/sequence/strength combinations.

**Supplementary Figure S10.**
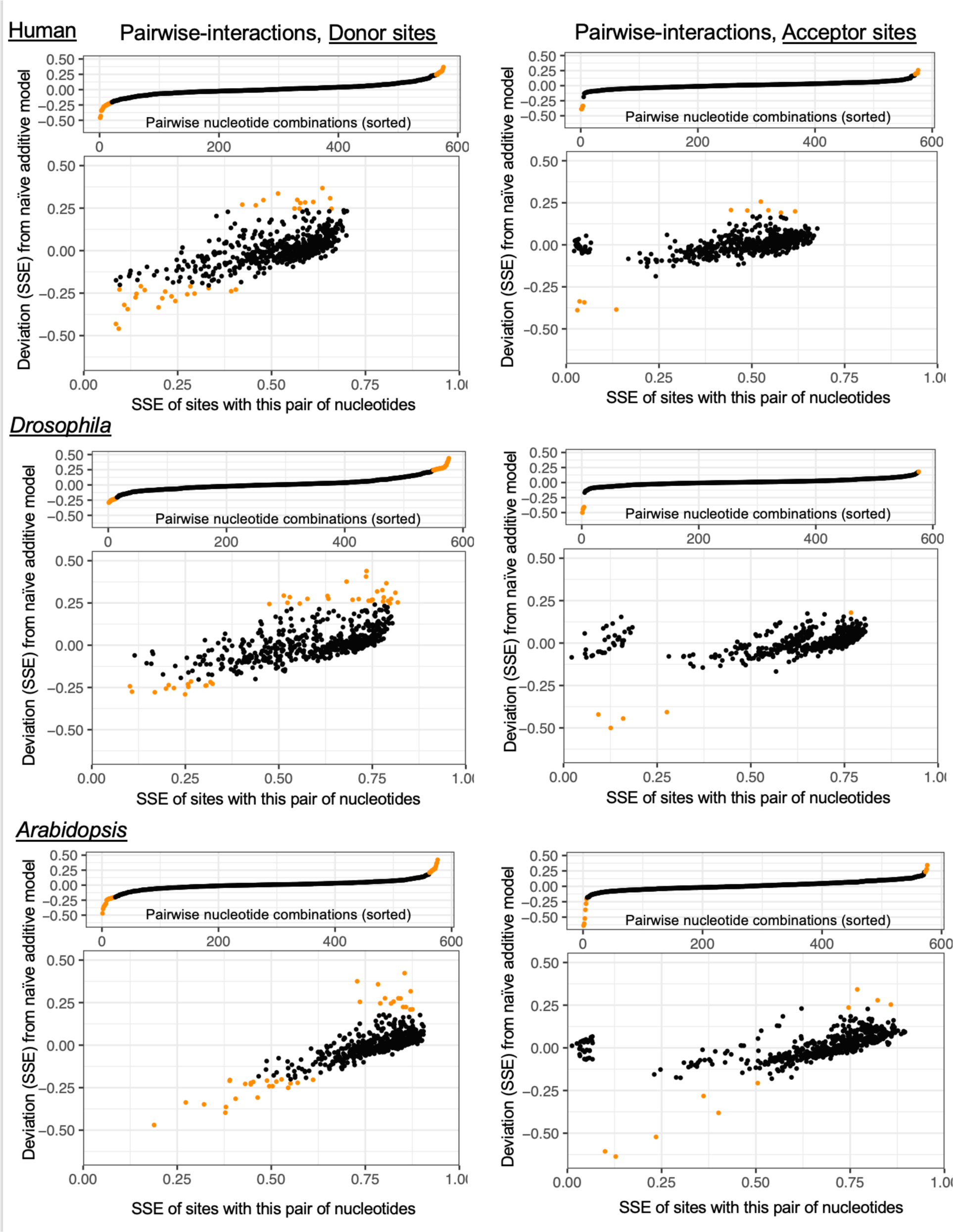
Pairwise combinations of nucleotides around the splice site whose Splice-site Strength Estimate (SSE) deviate from a naïve additive model. Left – Donor sites (GT only), Right - Acceptor sites (AG only). Each point represents a single pairwise combination of nucleotides around the splice site, -3 to +7 in donors, and -7 to +3 in acceptors. Upper panels – nucleotide pairs which deviate from the additive model are identified in orange. Lower panels – nucleotide pairs are plotted with their SSE (x-axis) and deviation from the additive model (y-axis) are identified in orange.

**Supplementary Figure S11.**
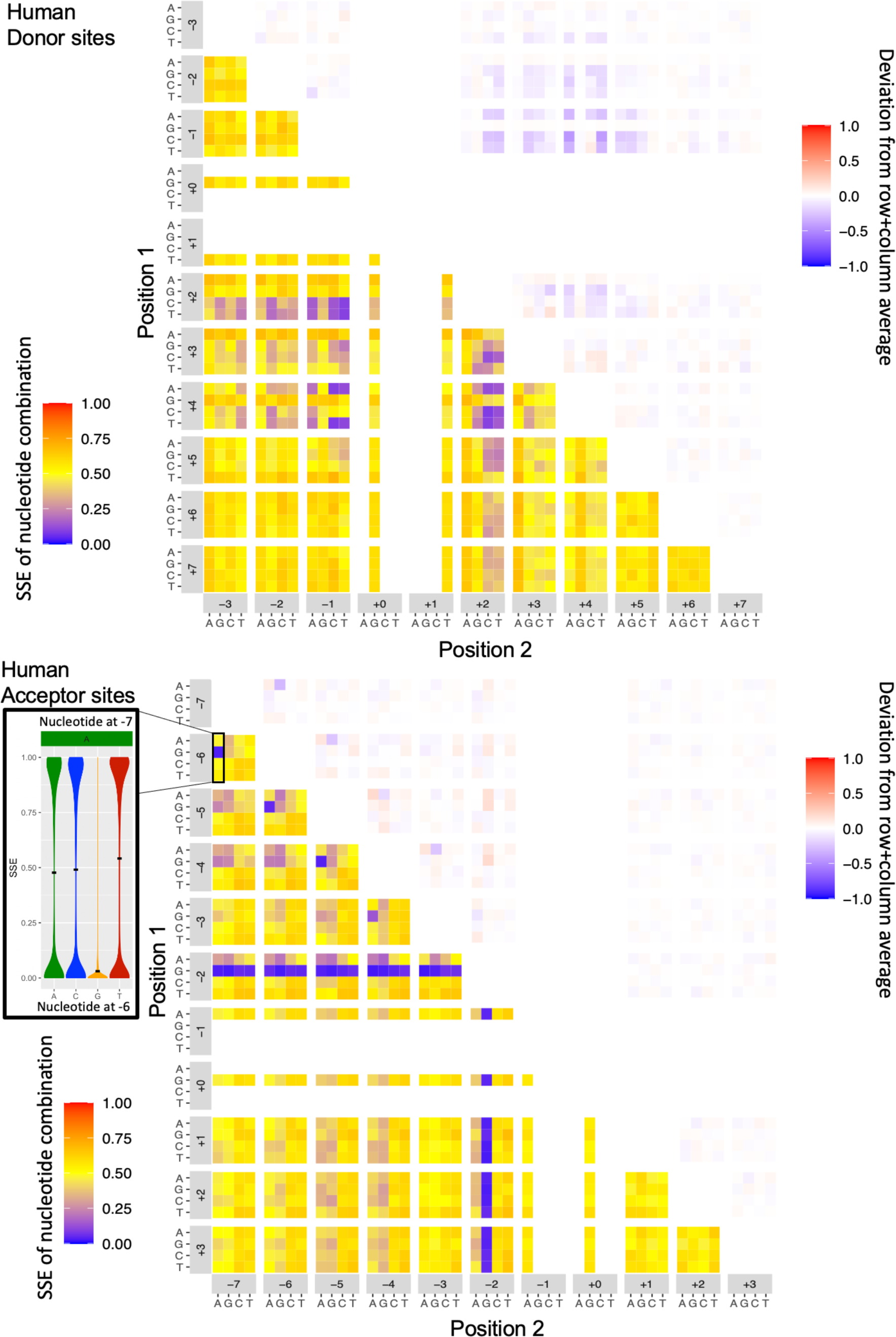
Heatmaps of Splice-site Strength Estimates (SSE) of pairwise combinations of nucleotides around human splice sites. Top – Donor sites (GT only), Bottom – Acceptor sites (AG only). Bottom left of plots show the SSE of the combination. Top right of plots show the deviation of these combinations from an additive model. Inset shows the SSE of nucleotides at the -6 position of splice acceptor sites which harbor an adenine at position -7.

**Supplementary Figure S12.**
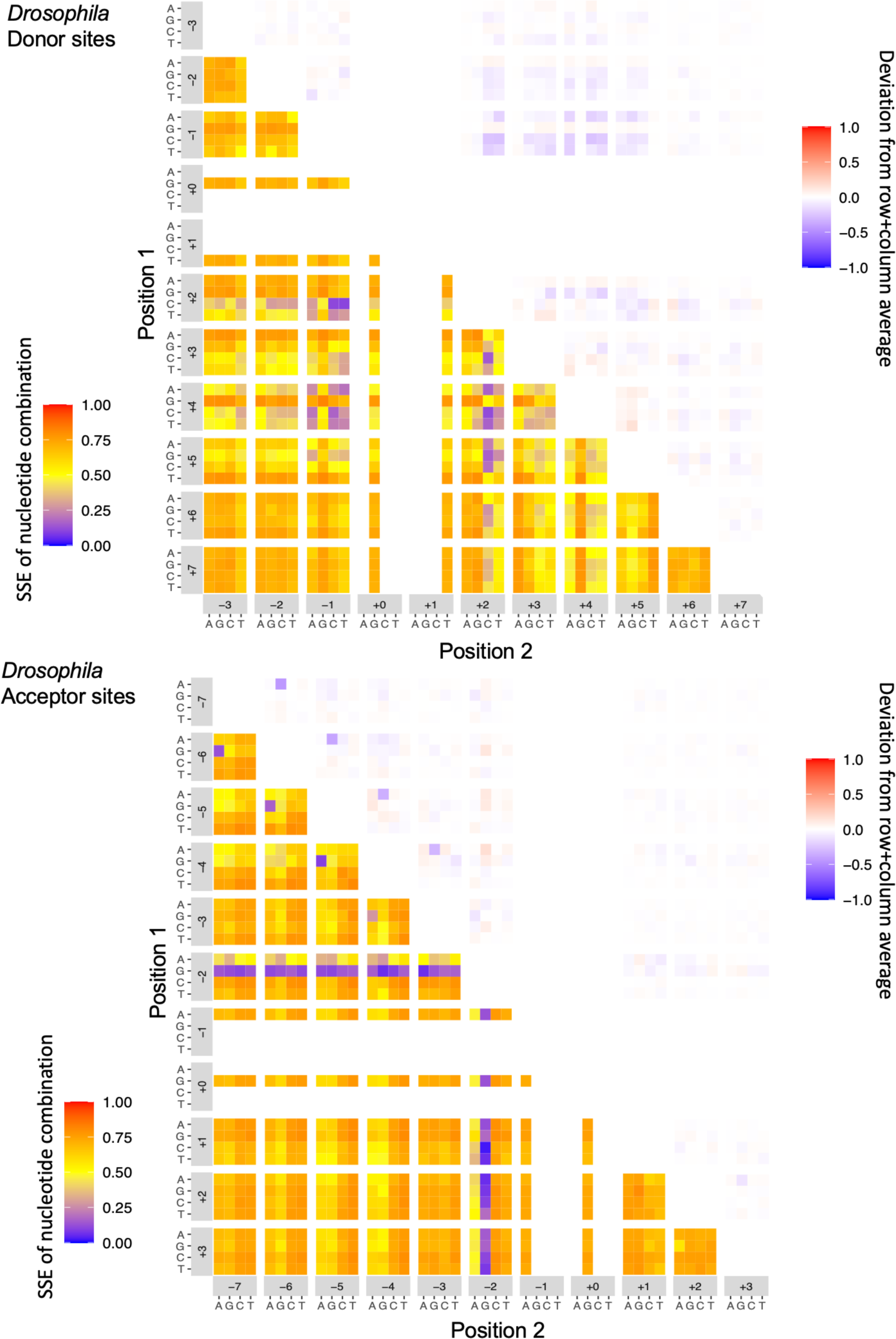
Heatmaps of Splice-site Strength Estimates (SSE) of pairwise combinations of nucleotides around Drosophila splice sites. Top – Donor sites (GT only), Bottom – Acceptor sites (AG only). Bottom left of plots show the SSE of the combination. Top right of plots show the deviation of these combinations from an additive model.

**Supplementary Figure S13.**
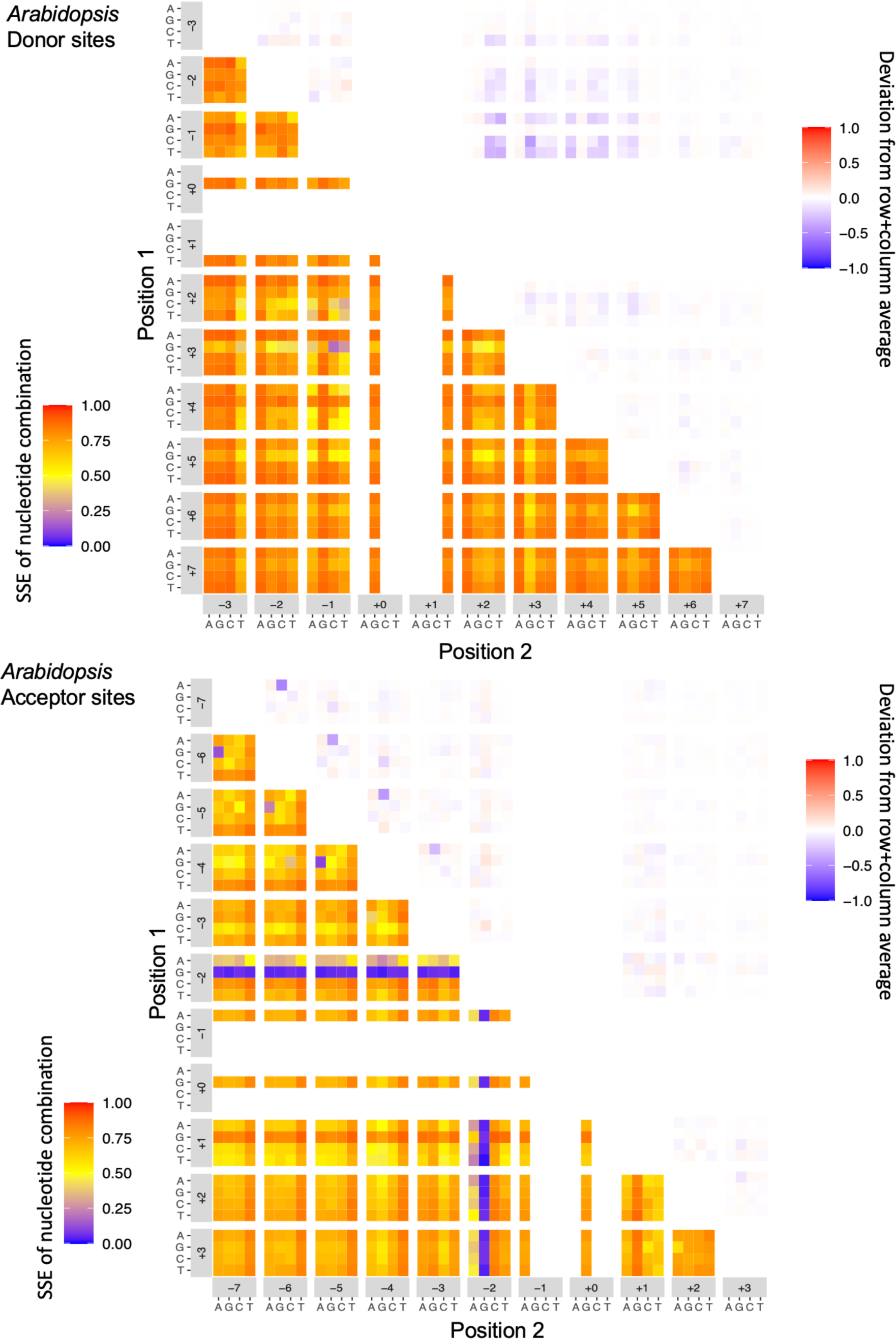
Heatmaps of Splice-site Strength Estimates (SSE) of pairwise combinations of nucleotides around Arabidopsis splice sites. Top – Donor sites (GT only), Bottom – Acceptor sites (AG only). Bottom left of plots show the SSE of the combination. Top right of plots show the deviation of these combinations from an additive model.

**Supplementary Figure S14.**
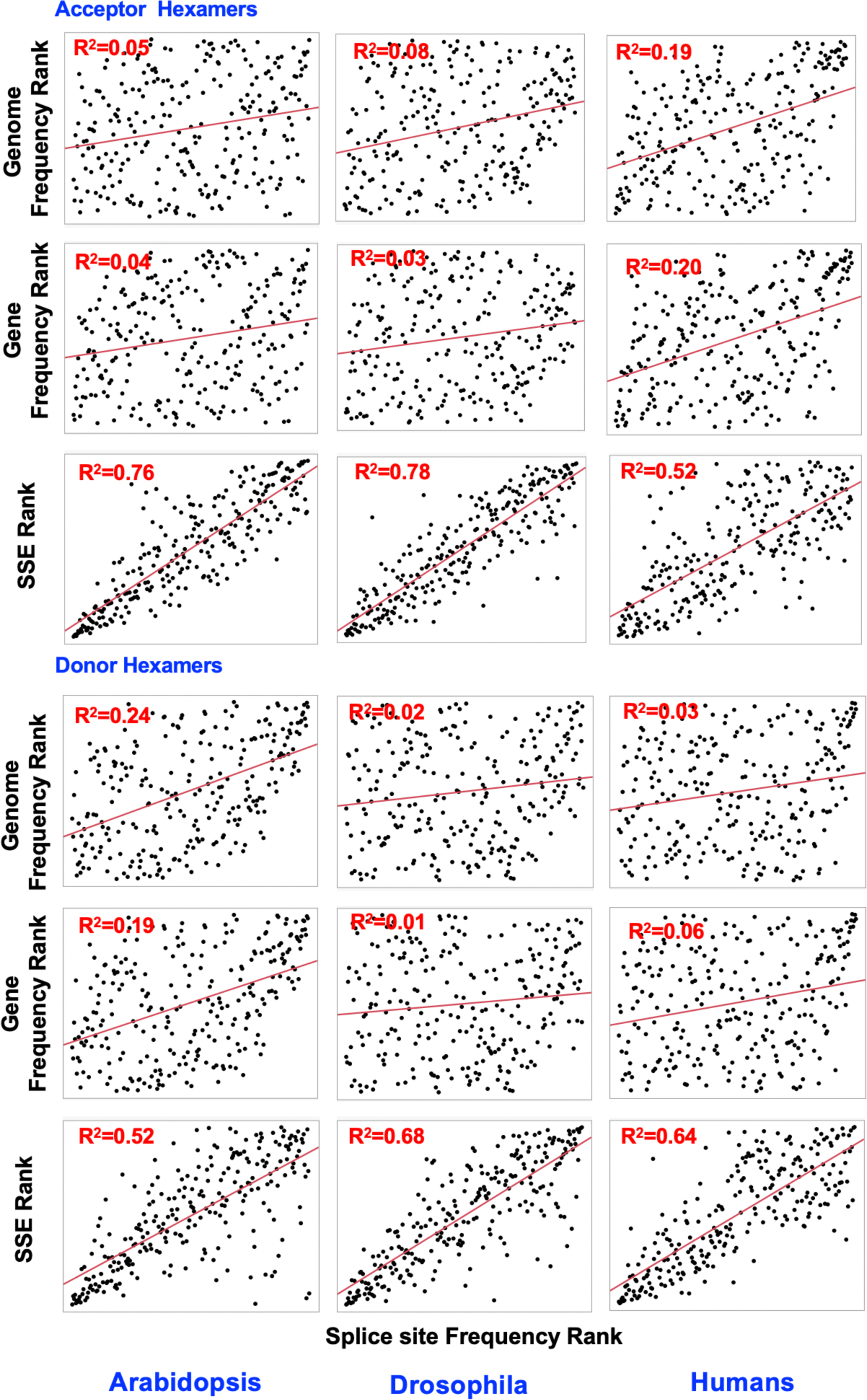
Hexamer frequency at the splice-sites is correlated with splice-site strength but not with the frequency of the hexamers in the genome or within protein coding genes. Hexamer frequency ranks are plotted against their corresponding ranks in the genome or genes, or the ranking of the hexamers based on splice-site strength estimates. R^2^ values are indicated on top for each of the correlations.

**Supplementary Figure S15.**
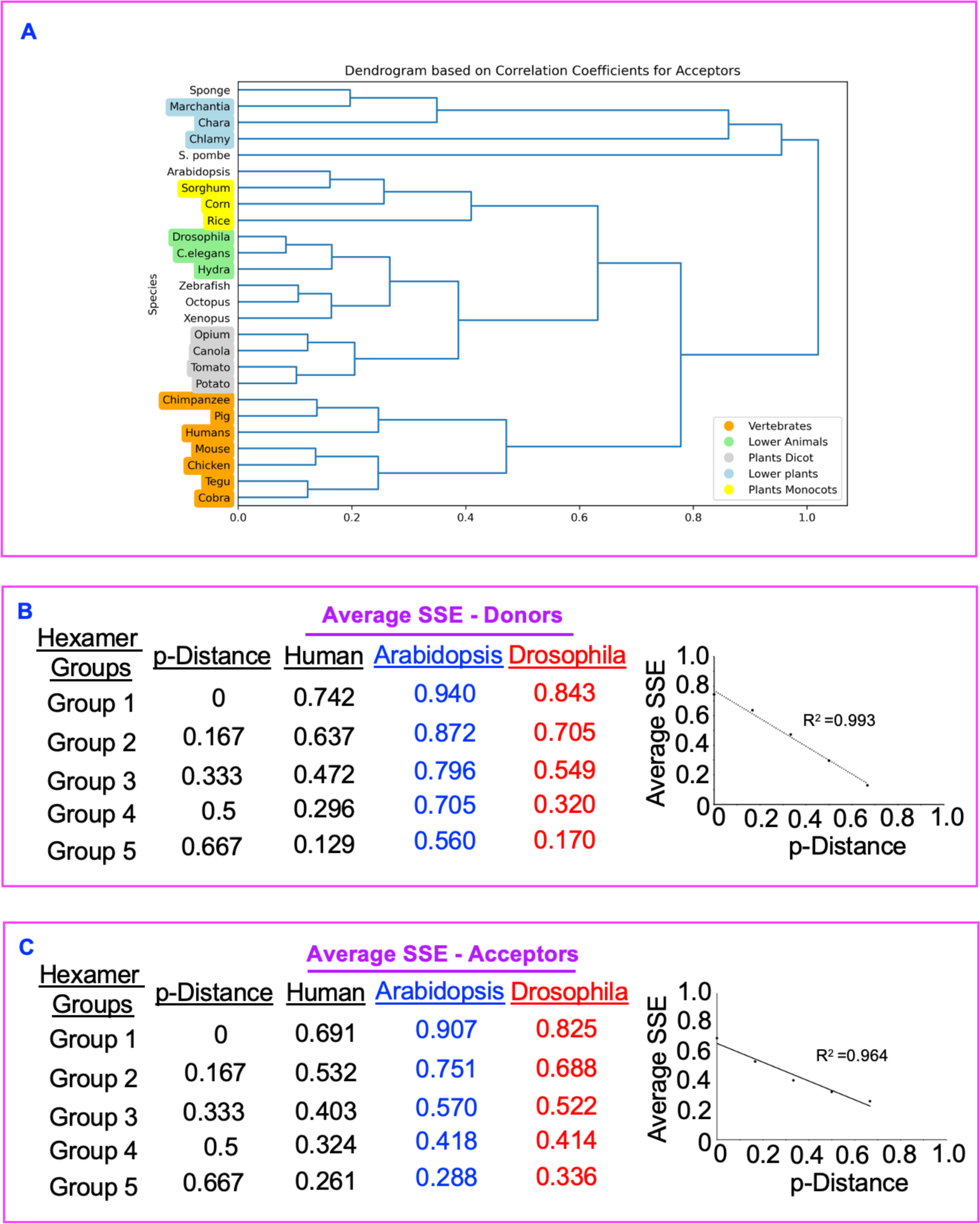
Hexamer effect on donors could be accounted for by variation in U1 snRNA base pairing for donor sites (strongest hexamer for donors) and sequence distance from the strongest hexamer for acceptors. A) A dendrogram of the rank correlations based on the hexamers surrounding acceptors in multiple species groups related species together. B-C) Sequence distance-based grouping of hexamers and their average strengths for donors (B) and acceptors (c) exhibits a near perfect correlation in all three species. Only the R^2^ values for human are shown and other species displayed similar high correlations.

**Supplementary Figure S16.**
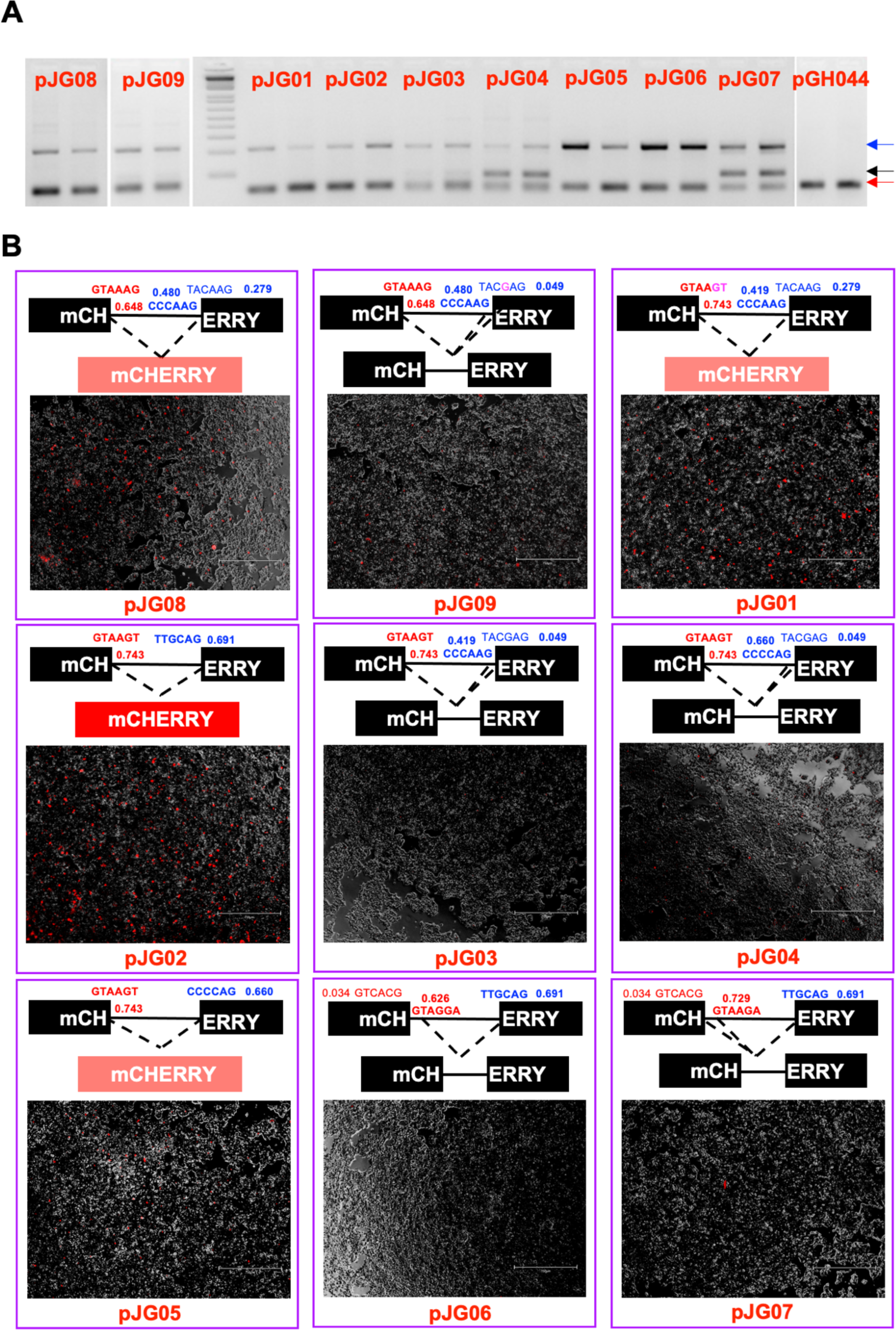
Hexamer variations confer differential splicing that is consistent with hexamer rank order. A) RT-PCR analysis of splicing with different hexamer sequence variants in the *MYO15B* gene. B) mCHERRY mini gene assays with *MYO15B* intron with varied hexamer combinations. The specific mutations are shown in purple, and the donor and acceptors are shown in red and blue, respectively.

**Supplementary Figure S17.**
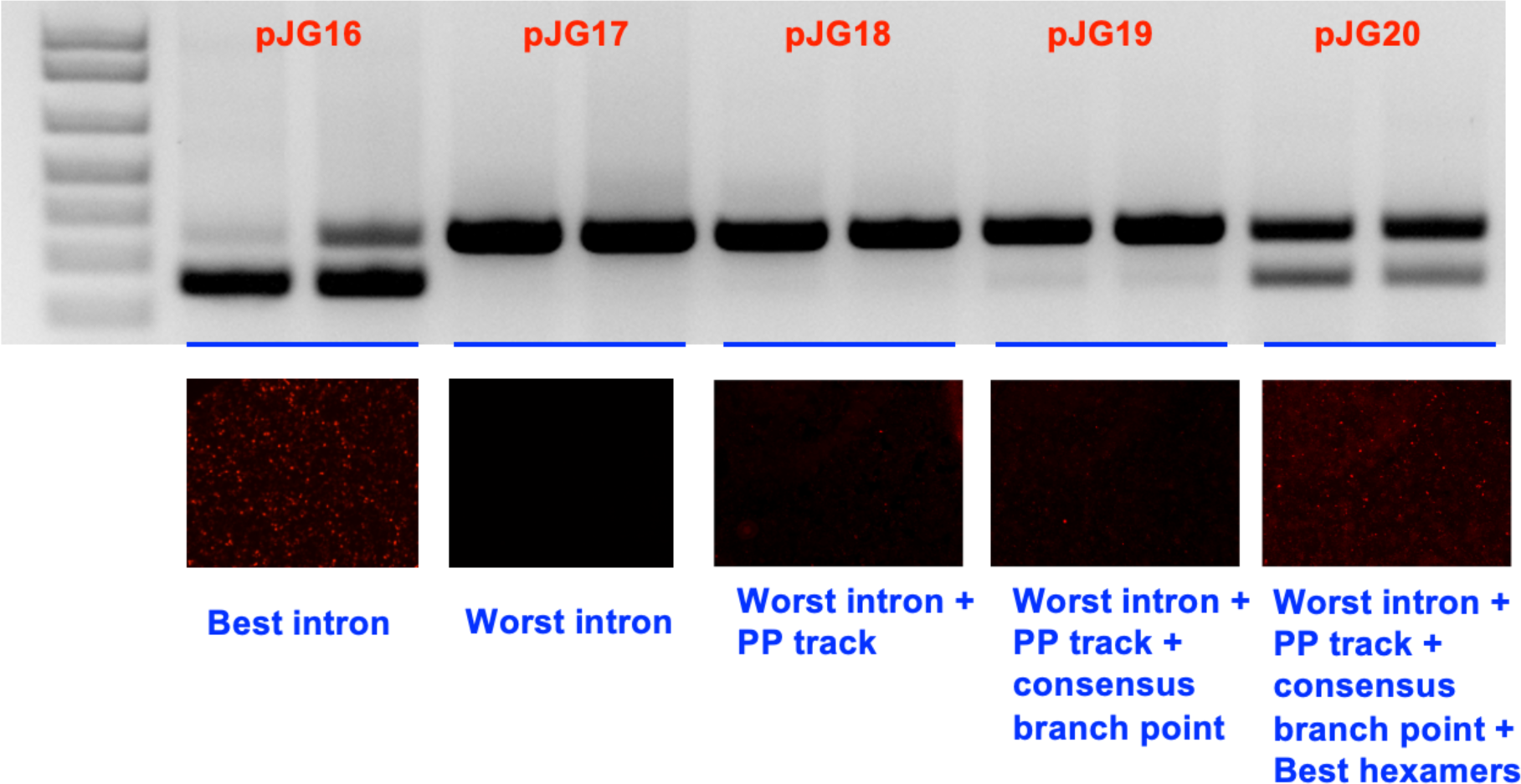
Hexamers make a significant difference to splicing. An RT-PCR analysis of the best/worst synthetic introns along with modified constructs harbouring PP tract, branch point and the hexamers.

**Supplementary Table S1. The number and percentage of splice-sites that show either 20% or 50% difference between individuals in their usage in Arabidopsis, Drosophila and Humans.**

**Supplementary Table S2. SpliSER-GWAS results for all “GWASable” splice-site mutations in Arabidopsis, Drosophila and Humans.**

**Supplementary Table S3. SpliSER-GWAS results for all associations detected in *Arabidopsis thaliana.***

**Supplementary Table S4. SpliSER-GWAS results for all associations detected in *Drosophila melanogaster.***

**Supplementary Table S5. SpliSER-GWAS results for all associations detected in the Humans.**

**Supplementary Table S6. Inferred best and worst nucleotides from the associated SNPs that are also closest to the splice-sites from GWAS data. Data for Arabidopsis, Drosophila and Humans and a combined analysis of all three species for donors and acceptors is given. Top rows represent the positions and bottom row represents the inferred nucleotides.**

**Supplementary Table S7. Constructs generated to test introns with various sequence combinations in this study.**

**Supplementary Table S8. The mean Splice-site Strength Estimate (SSE) of splice donor sites, with or without particular nucleotides in the -3 to +7 region. Downsampled in each species to 1-2 million site/sequence/strength combinations.**

**Supplementary Table S9. The mean Splice-site Strength Estimate (SSE) of splice acceptor sites, with or without particular nucleotides in the -7 to +3 region. Downsampled in each species to 1-2 million site/sequence/strength combinations.**

**Supplementary Table S10. The mean Splice-site Strength Estimate (SSE) of splice donor sites, with pairwise combinations of nucleotides at different positions in the -3 to +7 region, and their deviation from an additive model. Downsampled in each species to 1-2 million site/sequence/strength combinations.**

**Supplementary Table S11. The mean Splice-site Strength Estimate (SSE) of splice acceptor sites, with pairwise combinations of nucleotides at different positions in the -7 to +3 region, and their deviation from an additive model. Downsampled in each species to 1-2 million site/sequence/strength combinations.**

**Supplementary Table S12. Hexamer ranking explains most of the splice-site choice across the genome in diverse organisms. Here, percentage explained refers to the number of splice-sites with highest average strength hexamer in a window of +/-50bp without applying a penalty for duplication.**

**Supplementary Table S13. Donor hexamer counts, strengths, and ranks for all species. *Sc -Saccharomyces cerevisiae* baker’s yeast, *Sp - Saccharomyces pombe* budding yeast, Algae – *Chlamydomonas*, *Chara* - Charaphycean algae, *Mp - Marchantia polymorpha* liverwort, *Os - Oryzae sativa* rice, *Zm – Zea mays* corn, Sorghum – Sorghum bicolor, *AT- Arabidopsis thaliana*, Canola – *Brassica napus*, Opium – Opium, Potato – potato, *Sl – Solanum lycopersicum* tomato, Sponge – sponges, *Hv – Hydra vulgaris*, *Ce – Caenorhabditis elegans* worm, *Dm – Drosophila melanogaster* flies, Octopus – octopus, *Dr – Danio rario* Zebrafish, *Xenopus – Xenopus* frog, Cobra – cobra, Chicken – chicken, *Mm – Mus musculus* Mouse, Pig – pig, *Pt – Pan troglodytes* Chimpanzee, *Hs – Homo sapiens* humans. For Arabidopsis, Drosophila and Humans, the rankings were calculated from all individuals that were part of the GWAS data. For all other species, a single transcriptome data was used to generate the rankings.**

**Supplementary Table S14. Acceptor hexamer counts, strengths, and ranks for all species. *Sc -Saccharomyces cerevisiae* baker’s yeast, *Sp - Saccharomyces pombe* budding yeast, Algae – *Chlamydomonas*, *Chara* - Charaphycean algae, *Mp - Marchantia polymorpha* liverwort, *Os - Oryzae sativa* rice, *Zm – Zea mays* corn, Sorghum – Sorghum bicolor, *AT- Arabidopsis thaliana*, Canola – *Brassica napus*, Opium – Opium, Potato – potato, *Sl – Solanum lycopersicum* tomato, Sponge – sponges, *Hv – Hydra vulgaris*, *Ce – Caenorhabditis elegans* worm, *Dm – Drosophila melanogaster* flies, Octopus – octopus, *Dr – Danio rario* Zebrafish, *Xenopus – Xenopus* frog, Cobra – cobra, Chicken – chicken, *Mm – Mus musculus* Mouse, Pig – pig, *Pt – Pan troglodytes* Chimpanzee, *Hs – Homo sapiens* humans. For Arabidopsis, Drosophila and Humans, the rankings were calculated from all individuals that were part of the GWAS data. For all other species, a single transcriptome data was used to generate the rankings.**

**Supplementary Table S15. R^2^ values of correlations between hexamer rankings of acceptors (A & B) and donors (C & D) based on counts (A & C) or splice-site strength (B & D) between diverse species.**

**Supplementary Table S16. PhenoScanner scan results for phenotypes associated with SNPs that are detected to be the highest associated SNPs in the SpliSER-GWAS analysis of the human heart atrial tissue RNA-seq data.**

